# A contribution to the anatomy of two rare cetacean species: the hourglass dolphin (*Lagenorhynchus cruciger*) and the spectacled porpoise (*Phocoena dioptrica*)

**DOI:** 10.1101/2024.11.06.622215

**Authors:** Jean-Marie Graïc, Tommaso Gerussi, Bruno Cozzi, Rebecca M. Boys, Brian Chin Wing Kot, Matthew R. Perrott, Kane Fleury, Tabris Yik To Chung, Henry Chun Lok Tsui, Emma Burns, Trudi Webster, Stuart Hunter, Emma L. Betty, Odette Howarth, Carolina Loch, Sophie White, Steve Dawson, William Rayment, Ros Cole, Derek Cox, Tom Waterhouse, Hannah Hendriks, Anton van Helden, Muriel Johnstone, Ramari Oliphant Stewart, R. Ewan Fordyce, Karen A. Stockin

## Abstract

The anatomical description of the hourglass dolphin (*Lagenorhynchus cruciger*) and the spectacled porpoise (*Phocoena dioptrica*) remains largely unexplored, due to limited specimen availability and preservation challenges. This study employed digital imaging techniques, conventional histology and computed tomography to provide visualisation of anatomical structures for a detailed analysis. We present a comprehensive analysis of the gross macroscopical and microscopical morphology of two hourglass dolphins and four spectacled porpoises. The hourglass dolphins were characterised by their distinctive black and white pigmentation and a hooked dorsal fin, while the spectacled porpoises were defined by their large dorsal fin, lack of a visible rostrum and unique eye markings. Morphometric measurements and skeletal characteristics aligned with the literature, while internal anatomy (organs and systems) were similar to other odontocetes. Although precise lung measurements were challenging, qualitative assessments indicated relatively large lungs for their body size, supporting the “short dive, big lung” hypothesis and suggesting that these species are not deep divers. The spectacled porpoise dorsal fin was uniquely large with a well-developed blood supply; this is hypothesised to act as a thermoregulatory window, helping to manage body heat. Overall, this study provides new data on the anatomy of the hourglass dolphin and spectacled porpoise, contributing insights that may influence future research on these rare species. The findings highlight the importance of anatomical studies in explaining evolutionary relationships within cetaceans and their ecological roles in the Southern Ocean ecosystems.

## Introduction

The hourglass dolphin (*Lagenorhynchus cruciger*, Quoy & Gaimard, 1824), and the spectacled porpoise (*Phocoena dioptrica*, Lahille, 1912) are two species of small (ca. 2 m) cetaceans that inhabit subantarctic and Antarctic waters (Fordyce et al., 1984; Brownell and Donahue, 1999; Hammond et al., 2008; Cipriano, 2018; Goodall and Brownell et al., 2018). While aspects of their external morphology have been reported (Cipriano, 2018; Goodall and Brownell et al., 2018; refer to Table 1), internal anatomy has seldom been considered, likely because of their southern, limited geographic range and correspondingly poor access to fresh, intact specimens. The hourglass dolphin belongs to the family *Delphinidae*, though its taxonomy is still under debate with a recent proposal to shift from *Lagenorhynchus* to *Sagimatus* genus (Vollmer et al., 2019; see Jefferson et al., 2015 for taxonomy and general description). Recent genomic analysis offers further insight to the hourglass dolphin and its placement within the *Delphininae* subfamily (McGrath et al., 2025). The most relevant external characteristics are the distinctive white and black pigmentation of the flanks and the markedly hooked dorsal fin (Figure 1A). The common name arises from the hourglass shaped white marking extending from the beak to the tail flukes (Jefferson et al., 2015; Cipriano, 2018).

**Figure 1:**
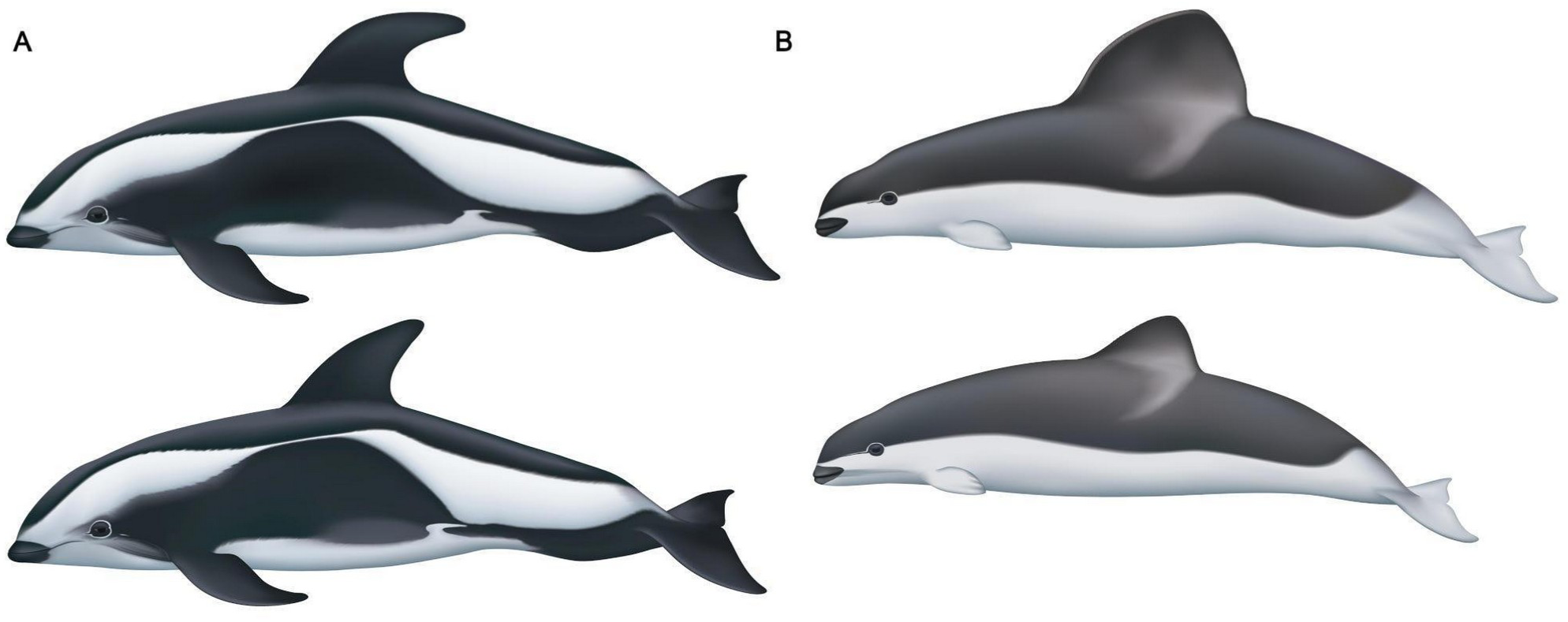
(A) Typical aspect of an adult male (top) and female (bottom) hourglass dolphin (*L. cruciger*) and (B) adult male (top) and female (bottom) spectacled porpoise (*P. dioptrica*). Illustrations kindly provided by Uko Gorter (https://ukogorter.com/).

**Table 1.** Selected data available on hourglass dolphin and spectacled porpoise specimens available in the published literature.

| Parameters | Hourglass dolphin |  | Spectacled porpoise |  | Reference |
| --- | --- | --- | --- | --- | --- |
|  | Value | n | Value | n |  |
| Total length (cm) | ♂ 162,6 - 187<br>♀ 142 - 182,9 | 6<br>7 | ♂ 109 - 224<br>♀ 119 - 203,5<br>? 94 - 101 | 8<br>13<br>11 | Goodall et al., 1997;<br>Brownell and Donahue, 1999;<br>Gazitúa et al., 1999;<br>Fernández et al., 2003;<br>Brownell, 1999;<br>Evans et al., 2001;<br>Pinedo, 2002 |
| Weight (kg) | ♂ 93 - 100<br>♀ 73,5 - 88 | 3<br>2 | mixed sex<br>1.6 (foetus) - 115 | 7 | Goodall et al., 1997;<br>Gazitúa et al., 1999;<br>Fernández et al., 2003;<br>Brownell, 1999;<br>Evans et al., 2001;<br>Pinedo, 2002 |
| Condylbasal length (mm) | 316 – 370 | 11 | 276 – 424 | 54 | Goodall et al. 1997<br>Gazitúa et al., 1999<br>Cipriano, 2018<br>Perrin et al., 2000 |
| Visible teeth (per side) | 26 – 34 top<br>27 – 35 bottom | 6 | 16 – 26 top<br>17 – 23 bottom |  | Perrin et al., 2000 |
| Vertebral column | C= 7<br>Th= 13 – 14<br>L= 18 – 19<br>Ca= 27+ | 9 | C= 7<br>Th= 14<br>L= 14 – 16<br>Ca= 32 – 33 |  | Marchesi et al., 2016<br>Perrin et al., 2000 |
| Ribs (per side) | 12 – 13 | 9 | 13 – 14 |  | Perrin et al., 2000 |
| Phalangeal formula | I= 2 – 3<br>II= 8 – 11<br>III= 6 – 8<br>IV= 2 – 4<br>V= 0 – 2 | 6 | I= 2<br>II= 7<br>III= 4<br>IV= 3<br>V= 4 |  | Perrin et al., 2000 |
| Intestine length | 18 – 19.7 m | 3 |  |  | Cipriano, 2018 |
| Number of reniculi | 670 (left kidney) | 1 |  |  | Cipriano, 2018 |
| Other particularities | Single vena cava,<br>absence of hepatic<br>sinus | 1 |  |  | Goodall, 1997;<br>Brownell, 1999 |

The spectacled porpoise belongs to the family *Phocoenidae*, and also suffers taxonomic uncertainty as to whether it is best placed within genus *Phocoena* or elsewhere (Jefferson et al., 2015). The species is characterised by a very large dorsal fin with a convex posterior margin, this feature is pronounced in males (Figure 1B). The eye is set within a small oval of black, with a thin dorsal semicircle of white, hence the common name “spectacled” (Jefferson et al., 2015; Goodall and Brownell, 2018).

While the literature describes the external morphology and skeleton of both species (Table 1), information on their visceral anatomy is scarce, possibly due to the limited number of specimens observed and to the decomposition status of the carcasses.

Our anatomical description of these little-known species yields new data that increase our understanding and may help in resolving their taxonomic status. Specifically, our study describes for the first time, the gross external morphology and the visceral macro- and micro-anatomy of six specimens (2 hourglass dolphin, 4 spectacled porpoise) examined postmortem.

## Materials and Methods

Conventional anatomical methods were applied, including dissection, photography, conventional histological staining, along with post-mortem computed tomography (PMCT). PMCT examination enhanced three-dimensional visualisation of organs and systems prior to dissection, allowing for volume calculations and providing comprehensive insights.

### Animal data

This study examined six specimens (n = 2 hourglass dolphins; n = 4 spectacled porpoise). All specimens originated from stranding events in New Zealand between 2010 and 2020 (Table 2). Each specimen was weighed, when possible, and measured prior to PMCT scanning.

**Table 2.** Specimen and strandings data of examined hourglass dolphins and spectacled porpoise examined in the study. Decomposition code according to the standards of Ijsseldijk et al., 2019.

| Species | Animal ID | Age class | Sex | Stranding date | Location stranding | Dissection date | Decomposition Code |
| --- | --- | --- | --- | --- | --- | --- | --- |
| hourglass dolphin | KS10-28Lc | Adult | M | 7 Sept 2010 | Flea Bay (43°86' S, 173°0' E), Akaroa, NZ | 10 Sept 2010 | 2-3 |
| hourglass dolphin | KS20-20Lc | Subadult | M | 5 Aug 2020 | Orepuki Beach (46°16' S, 167°43' E), Te Waewae Bay, NZ | 29 Sept 2020 | 1-2 |
| spectacled porpoise | KS14-45Pd/<br>X2020.76 | Adult | M | 17 Sept 2014 | Pipikaretu (45°80'S, 170°7'E), Otago Peninsula, NZ | 19 Sept 2014 | 1 |
| spectacled porpoise | KS14-37Pd/<br>X2020.77 | Subadult | M | 2 Oct 2014 | Caroline Bay Beach (44°38'S, 171°24'E), Timaru, NZ | 2 Oct 2014 | 1 |
| spectacled porpoise | KS15-29Pd/<br>VT3347 | Juvenile | F | 9 Aug 2015 | Bayleys Beach, (43°49'S, 172°36'E) Kaitorete Spit, Canterbury, NZ | 19 Aug 2015 | 3 |
| spectacled porpoise | KS20-07Pd | Adult | M | 14 Jan 2020 | Washdyke Lagoon, Canterbury, NZ | 22 Jan 2020 | 2 |

### PMCT scan data

PMCT scanning of the first hourglass dolphin (KS10-28Lc) was performed with a LightSpeed VCT CT scanner (GE Healthcare, USA), using the exposure parameters: 120 kV, 100 mA, 0.63 mm slice thickness, and sFOV of 45 cm. PMCT of the second hourglass (KS20-20Lc) was conducted using a CT 5000 Ingenuity CT scanner (Philips, Netherlands), with exposure parameters: 120 kV, 20 mA, 0.90 mm slice thickness, and sFOV of 50 cm. Three of the spectacled porpoises underwent PMCT scanning: KS14-37Pd/X2020.77 and KS14-45Pd/X2020.76 were conducted using a LightSpeed Pro 16 (GE Medical Systems); exposure parameters for KS14-45Pd/X2020.76 were 120 kV, 390 mA and 1.25 mm slice thickness, while for KS14-37Pd/X2020.77 were: 120 kV, 270 mA and 1.25 mm slice thickness. Due to logistical difficulties, the dorsal fin of KS14-45Pd/X2020.76 was scanned separately in the LightSpeed Pro 16 with scan parameters of 120 kV, 270 mA and slice thickness of 1.25 mm, this allowed for specific imaging of the dorsal fin structure. PMCT scanning of KS20-07Pd was conducted using an Optima CT660 (GE Medical Systems), with exposure parameters: 120 kV, 480 mA, and 0.63 mm slice thickness. Due to logistical difficulties the tail stock was removed and scanned separately in KS20-07Pd (exposure parameters remained the same), rendering it difficult to determine the precise number of lumbar vertebrae caudal to the dorsal fin. CT data from KS20-07Pd and KS20-20Lc cases was assessed using PMCT methodology (Kot et al., 2020; Granados-Zapata et al., 2022) and were used to guide the necropsy, thus improving findings of the conventional necropsy. Both scans were viewed using the TeraRecon iNtuition workstation (TeraRecon, San Mateo, CA, USA). Morphological and volumetric examinations were performed using Slicer 3D (https://www.slicer.org/).

### Dissection

Dissections occurred at the Cetacean Pathology Unit, Massey University Auckland (KS10-28Lc, KS20-20Lc, KS20-07Pd), AgResearch Invermay Campus, Dunedin (KS14-37Pd/X2020.77) and Tūhura Otago Museum, Dunedin (KS14-45Pd/X2020.76, KS15-29Pd/VT3347), New Zealand, following standardised sampling techniques (Geraci and Lounsbury 2007; IJisseldik et al., 2019). A more restricted examination was performed on the first hourglass dolphin (KS10-28Lc, no ingoa (cultural) name assigned) in order to preserve the integrity of the cadaver for cultural display. However, a full system dissection for KS20-20Lc (ingoa name, “Harua-tai-nui”) and all spectacled porpoises was permitted. Standardised histological tissue samples of key organs were fixed in 10% buffered formalin solution then trimmed, paraffin embedded and subsequently sectioned (8 μm) on a rotary microtome. Sections were stained with conventional hematoxylin-eosin stain, or with Masson’s trichrome.

The pectoral fins were analysed morphometrically, following Benke (1993). The condylo-basal length was also measured following Mead and Potter (1995).

## Results

### External appearance

Weight and morphometric measurements of each specimen are reported below in Table 3.

**Table 3.**
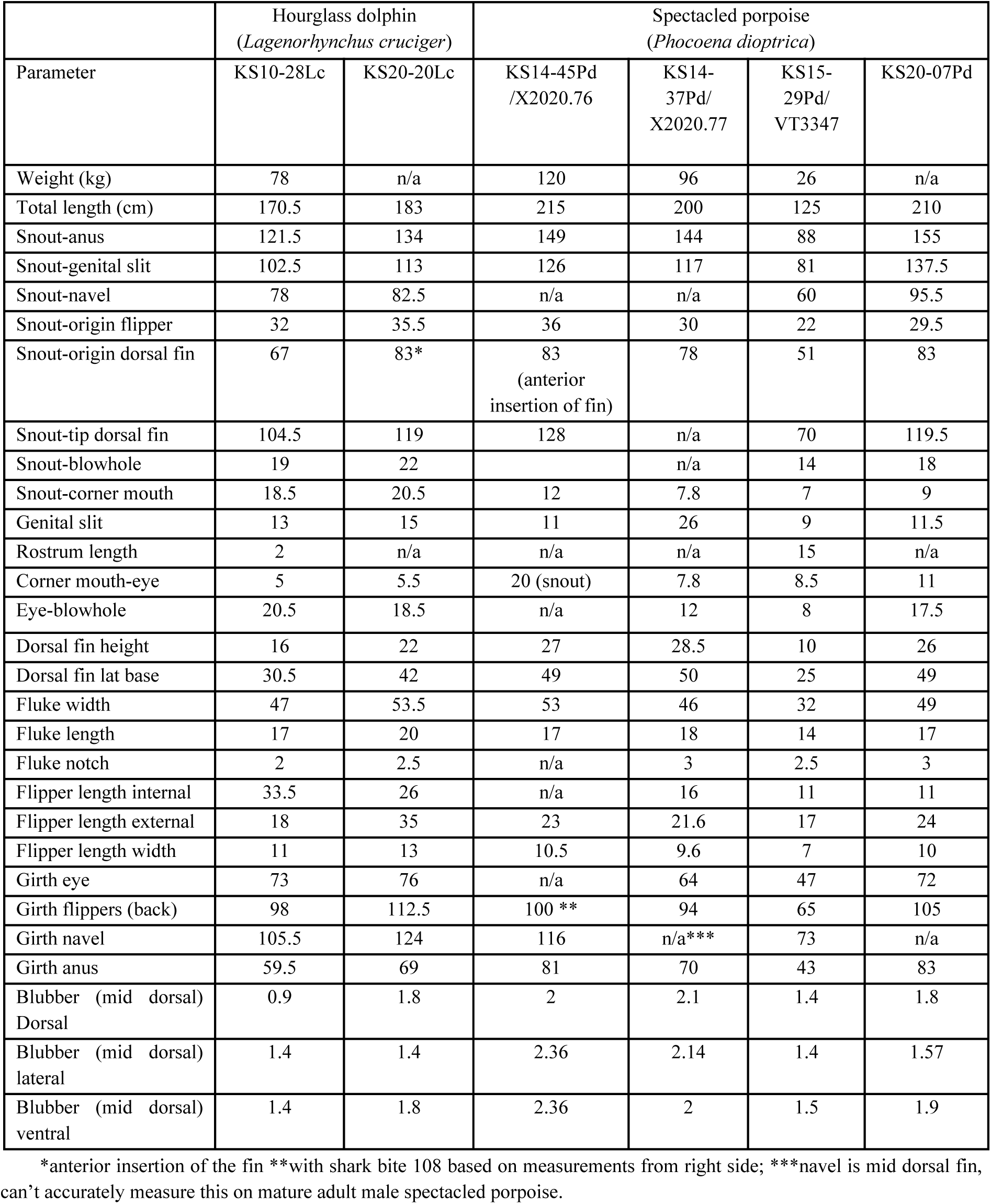
Weight (kg) and morphometrics (cm) of examined specimens.

The hourglass dolphins had a body shape broadly resembling a robust *Lagenorhynchus*, with a smoothly rounded head, and no protruding rostrum. The dorsal fin had a long base, was falcate, and notably tapered caudally, ending with a bluntly pointed tip. The pectoral fins displayed a narrower base and a notch on the caudal border adjoining the body. The caudal fin, or fluke, was relatively wide and thin. The caudal peduncle had an obvious ventral keel (Figure 2B), composed of dense connective tissue and blubber, with no additional musculature (Figure 2B). The characteristic colour pattern was similar to that described in the literature (see Figures 1 and 2). In common with the spectacled porpoise, the hourglass dolphins had an incomplete white line dividing the black eye spot with the black pattern of the head (Figure 2A).

**Figure 2:**
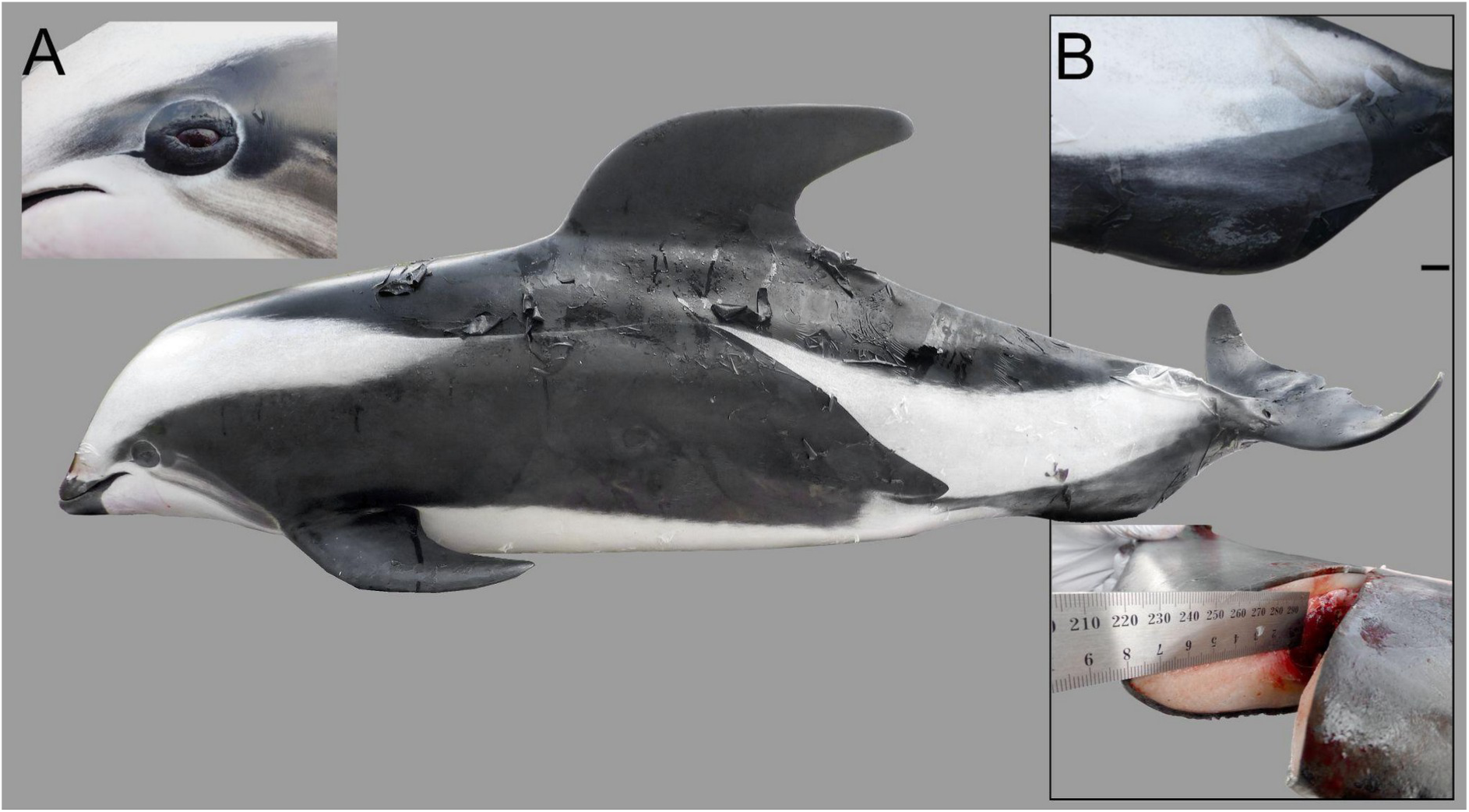
Adult male specimen of hourglass dolphin KS10-28Lc. Note: Pigmentation pattern with the characteristic shape of an hourglass. Note (A) white line around the eye spot and (B), caudal keel showing presence of connective tissue and absence of any musculature.

The spectacled porpoises showed a typical porpoise-like body form, having a squat and tapered body, a clear division between dorsal black and ventral white, rounded pectoral fins and a distinctively large dorsal fin with a convex trailing edge. The white line around the eye spot was incomplete (Figure 3).

**Figure 3:**
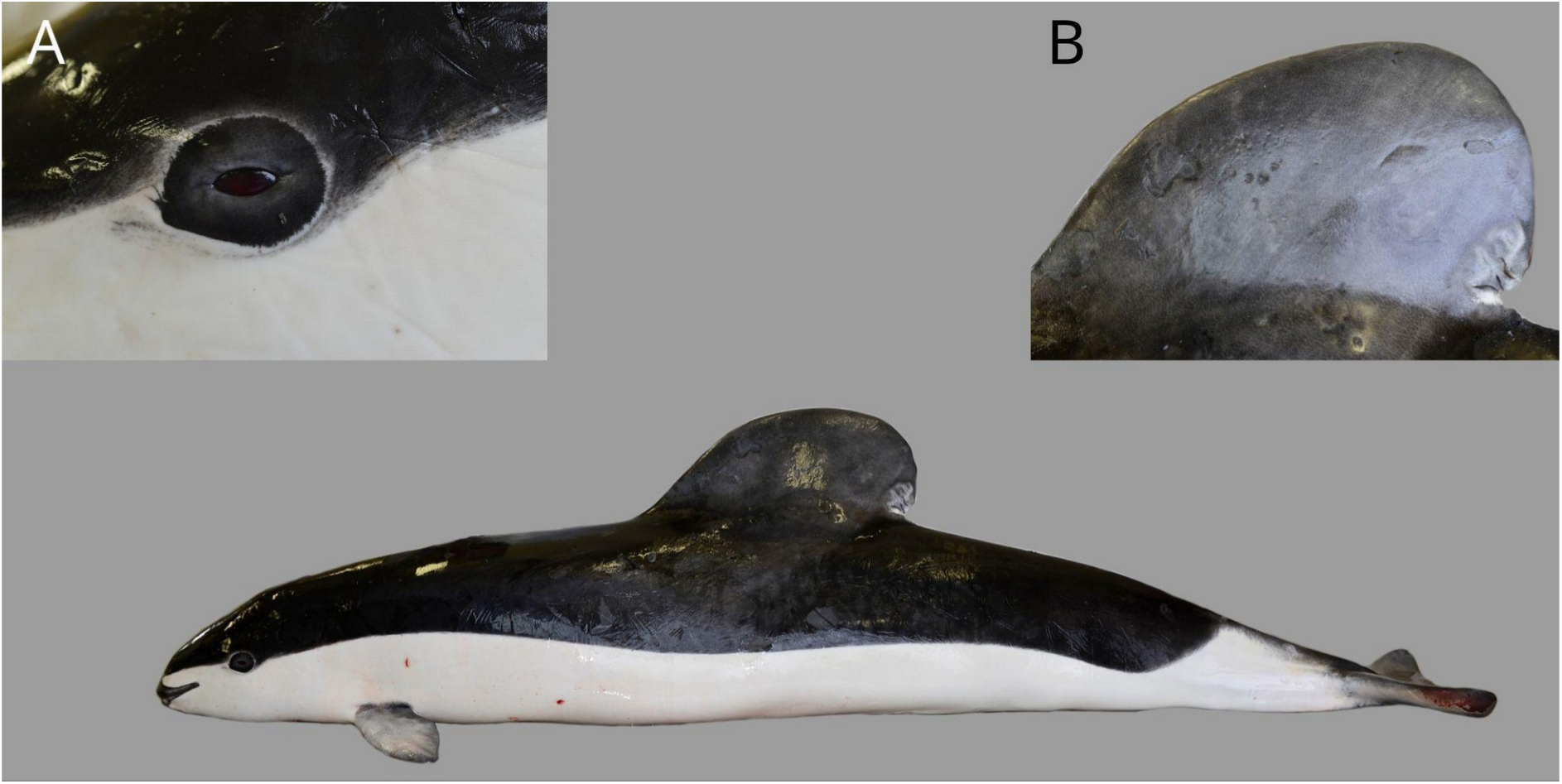
Adult male spectacled porpoise KS14-45Pd/X2020.76. Note (A), distinctive white rim around the black eye patch and (B), exaggerated, wide based dorsal fin.

In both hourglass dolphins, though not observed in spectacled porpoises, we detected four small pits in the skin of the rostrum, ca. 1 cm apart, likely representing remnants of the vibrissae.

### Osteology

In the hourglass dolphin, the two first cervical vertebrae (C1 and C2) were fused, while in the spectacled porpoise, C1 to C6 were fused (Figure 4). In both species, the atlas was large and relatively flat, while the remaining six vertebrae were tightly aligned with their body joined together. For vertebral formulae and details, see Table 4.

**Figure 4.**
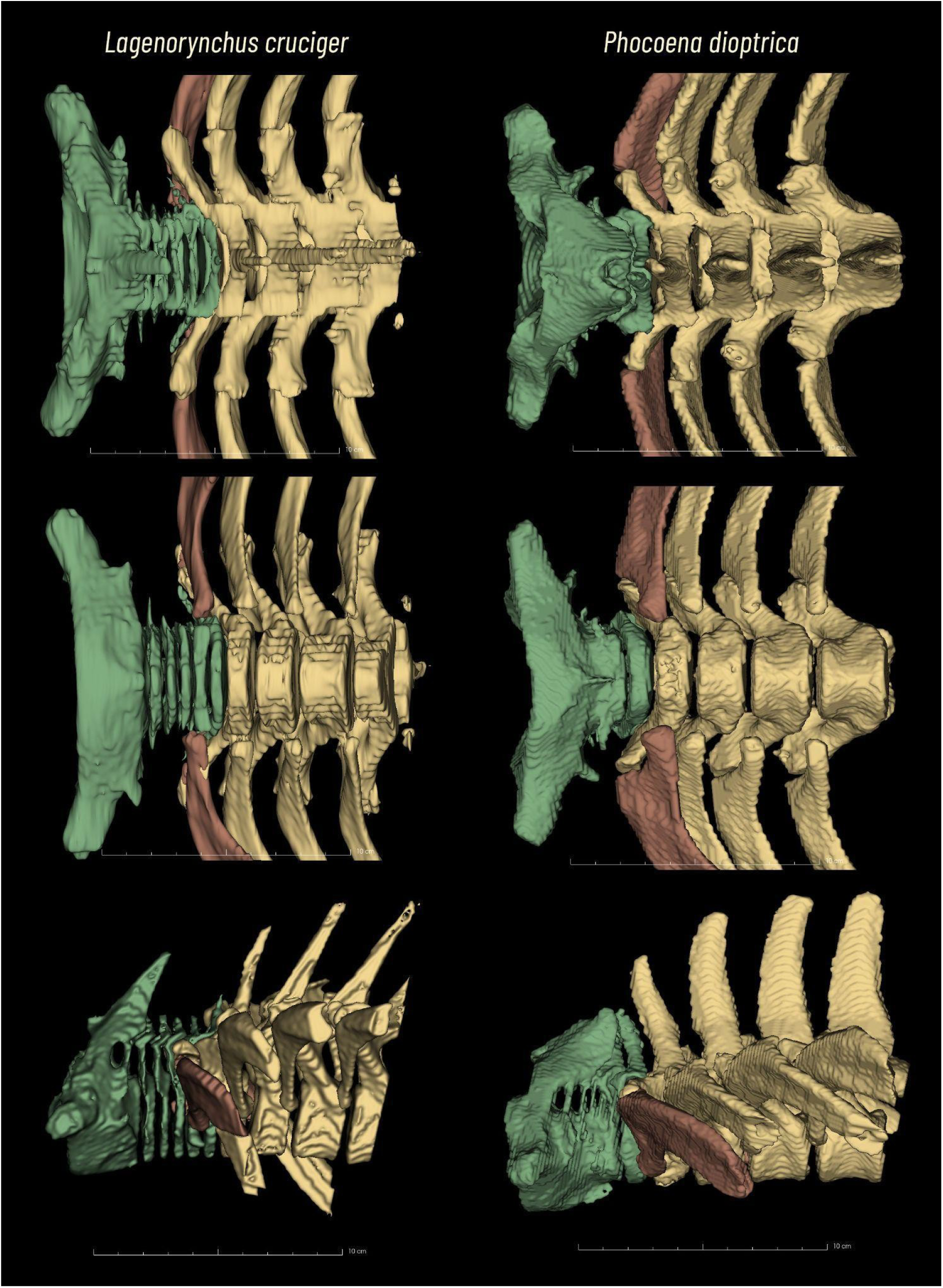
Adult hourglass dolphin KS20-20Lc and spectacled porpoise KS14-45Pd/X2020.76 cervical vertebrae segmented. In green C1-2, C3, C4, C5, C6 and C7; in yellow the first 4 thoracic vertebrae and ribs. In red, the first rib.

**Table 4.** Number of vertebrae per section, number of ribs and phalangeal formula (first number is left hand, second number is the right hand) by specimen. Note: CB = condylo-basal.

|  | Hourglass dolphin<br>(*Lagenorhynchus cruciger*) |  | Spectacled porpoise<br>(*Phocoena dioptrica*) |  |  |  |
| --- | --- | --- | --- | --- | --- | --- |
|  | KS10-28Lc | KS20-20Lc | KS14-45Pd<br>/X2020.76 | KS14-37Pd/<br>X2020.77 | KS15-29Pd/<br>VT3347 | KS20-07Pd |
| CB length | 319.6 | 362.8 | 307.2 | 293.4 | 239 | 289.6 |
| Cervical | 7 | 7 | 7 | n/a | 7 | 7 |
| Thoracic | 13 | 13 | 13 | n/a | 13 | 13 |
| Lumbar | 15 | 19 | na | n/a | 17 | n.d. |
| Coccygeal | nd | 16+20 | 32 | n/a | 37 | 27 |
| Ribs | 13 | 13 | 13 | n/a |  | 15? |
| Phalangeal<br>formula | I=<br>II=<br>III=<br>IV=<br>V= | I= 3; 2<br>II= 10; 10<br>III= 7; 7<br>IV= 3; 3<br>V= 3; 3 | I= 1; 1<br>II= 6<br>III= 5; 4<br>IV= 3; 3<br>V= 1; 1 | n/a | Right only<br>I=1<br>II= 6<br>III=5<br>IV=4<br>V=1 | (only R)<br>I= 2<br>II= 6<br>III= 5<br>IV= 4<br>V= 1 |

Both hourglass dolphin specimens had 13 thoracic vertebrae (Th), and an analogous number of ribs. Ribs 1 to 5 articulated with the sternum. The distal extremities of ribs 6 to 8 joined the distal bony aspect of the ribs rostral to them, while ribs 9 to 13 did not connect to the sternum. Ribs 1 to 6 articulated both with the transverse process of their vertebrae and the vertebral body or the vertebrae cranial to it (unlike in other mammals, in which it articulates between vertebrae), while ribs 7 to 13 articulated only with the transverse process of their vertebrae. In the spectacled porpoises, ribs 1 to 4 were connected to the sternum, ribs 5 to 8 were connected to the ribs rostral to them and the remaining ones were free of any relationship to the sternum (Figure 5).

**Figure 5:**
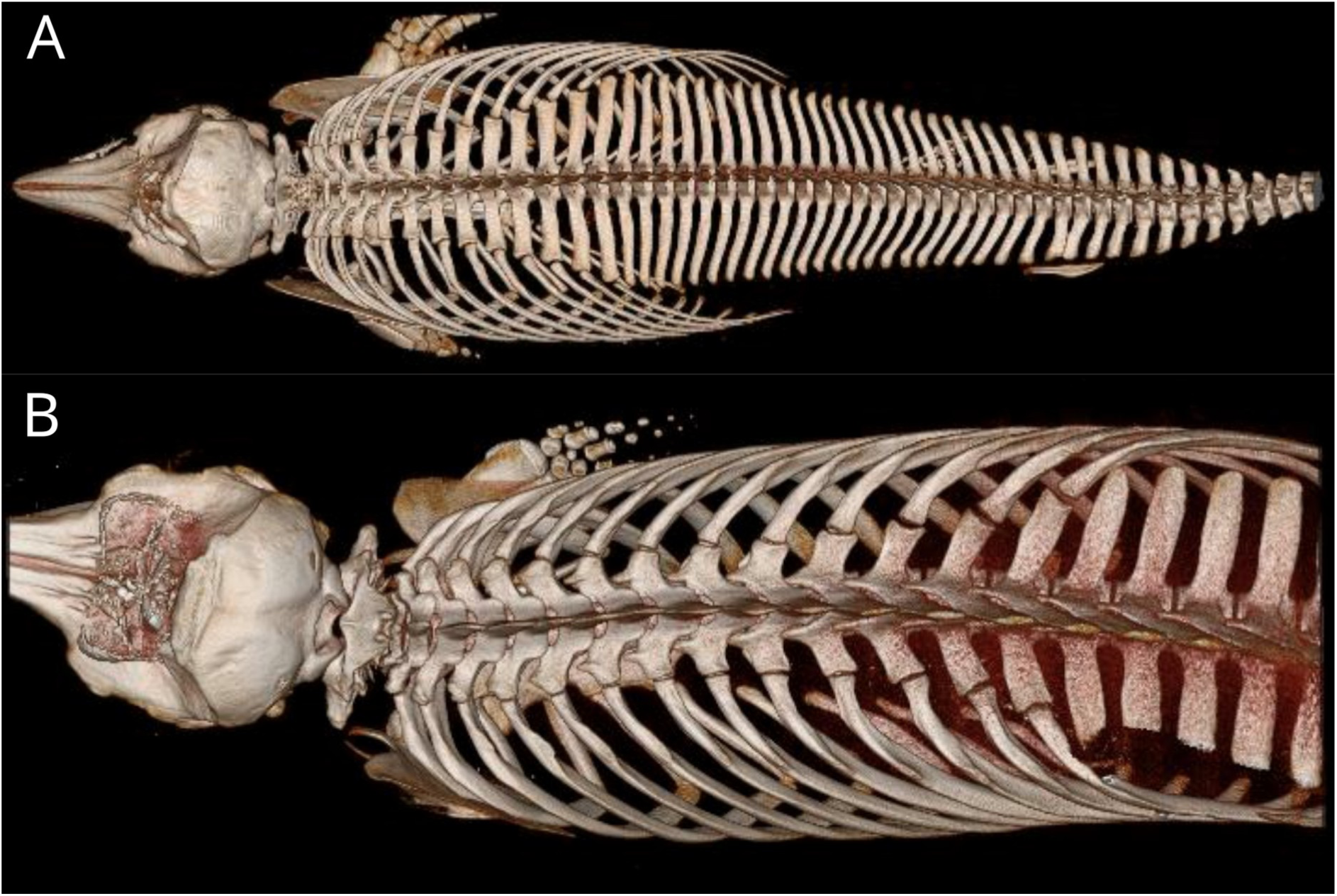
Representation of the skeleton up to the (A) caudal vertebrae of hourglass dolphin KS20- 20Lc and (B) lumbar vertebrae of spectacled porpoise KS20-07Pd, as displayed by 3D rendering of the CT scans.

The hyoid complex in hourglass dolphin KS20-20Lc displayed a typical *Lagenorhynchus* basihyoid connected to the stylohyoids, as described by Yablokov et al. (1974). In the spectacled porpoises, the basihyoid and thyrohyoid were fused in a porpoise-like unique flat bone, articulated cranially with the two stylohyoids which in turn, run back caudally (Figure 6).

**Figure 6:**
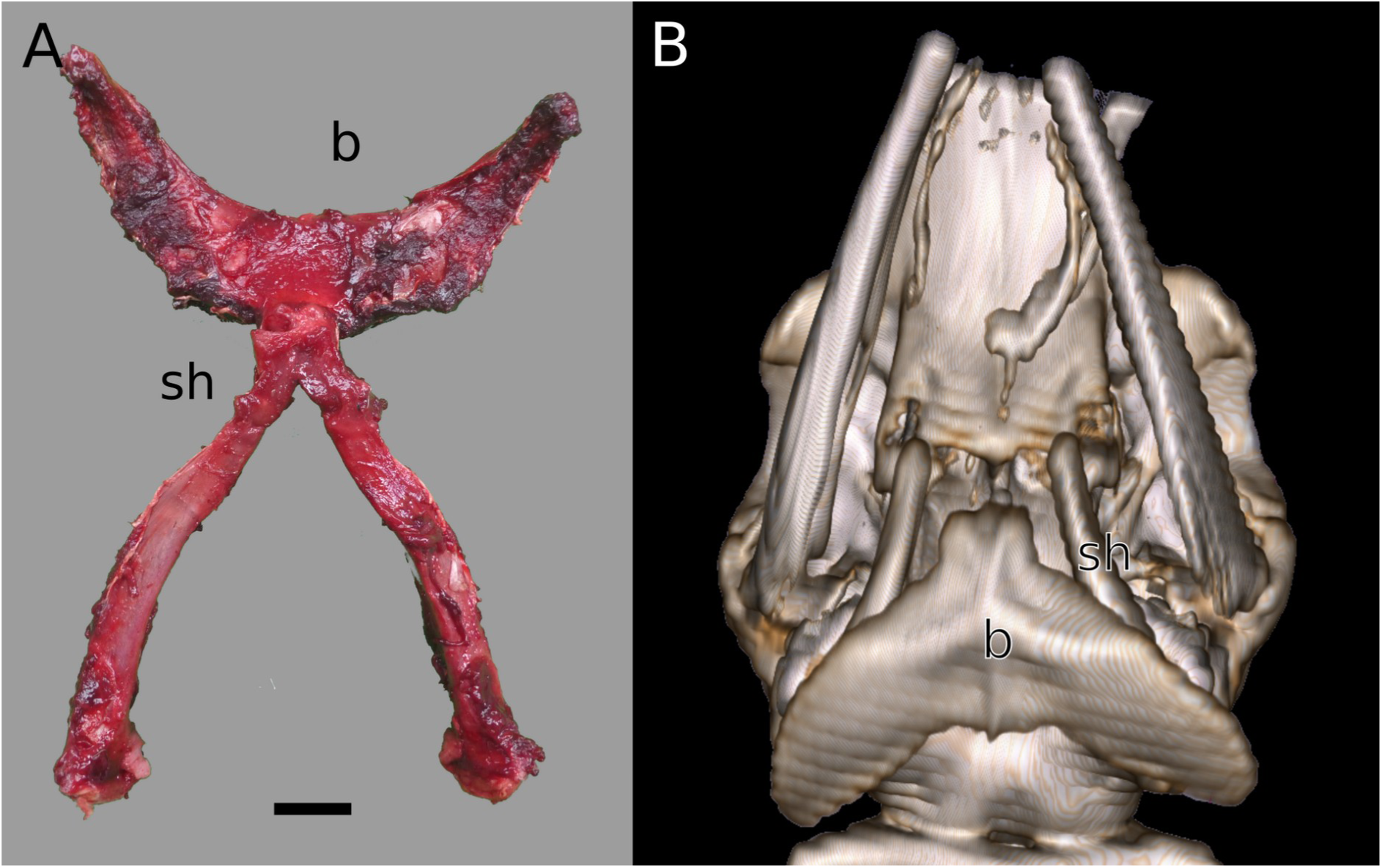
(A) Visualisation of the hyoid apparatus in a dorsal view of hourglass KS20-20Lc and (B) 3D reconstruction of the skeleton in a ventral view including the hyoid bone of spectacled porpoise KS20-07Pd. Note: b, basihyoid; sh, stylohyoid. Scale bar = 5 cm.

Pectoral morphometry was detailed in Figure 7 and Table 5.

**Figure 7:**
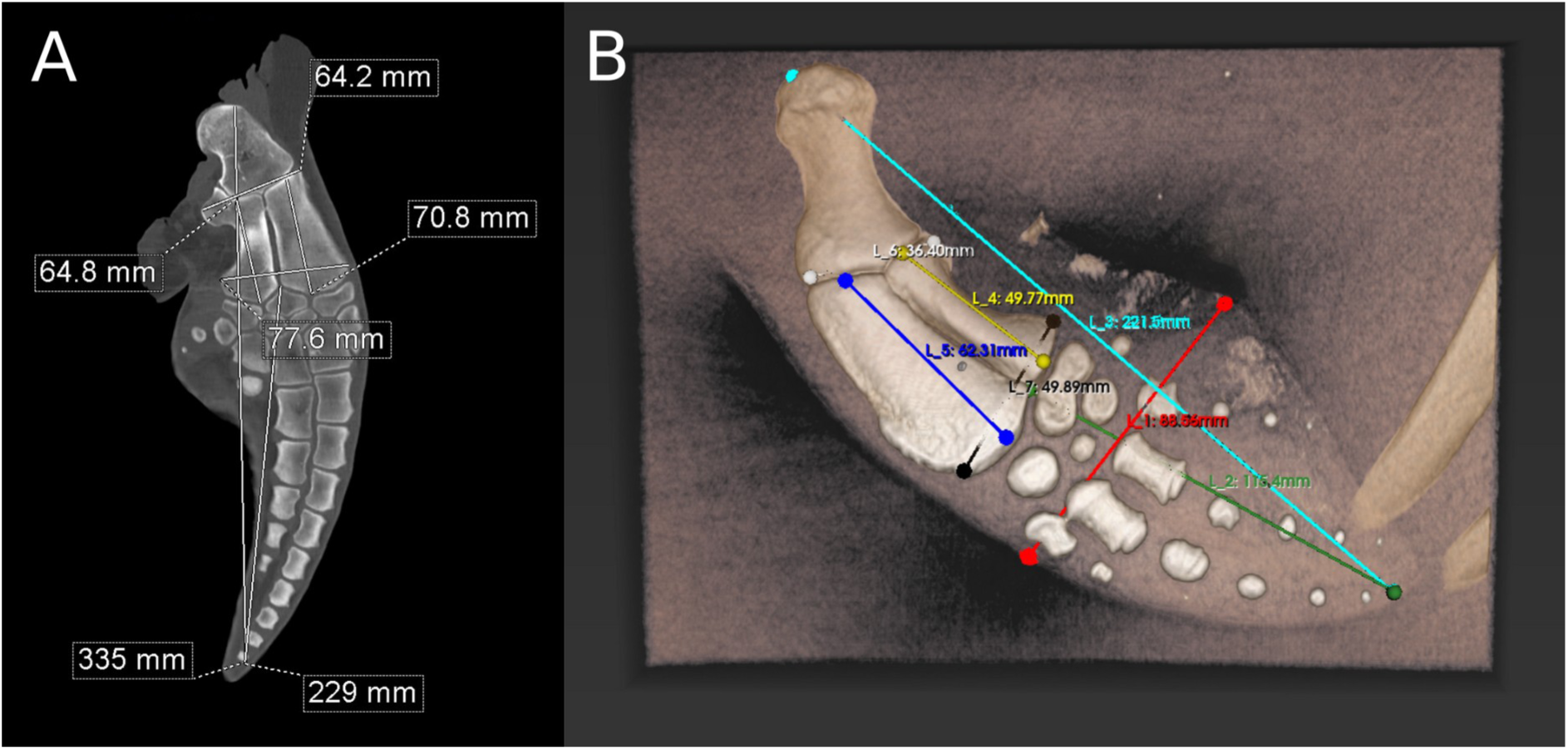
Visualisation of the skeletal characteristics with measurements (A) CT of left pectoral flipper of hourglass dolphin KS20-20Lc and (B) 3D reconstruction of the left pectoral flipper of KS20-07Pd with principal measurements.

**Table 5.** Morphometry of the pectoral fins.

| Measurement (cm) | Hourglass dolphin<br>(*Lagenorhynchus cruciger*) |  | Reference<br>(*Tursiops truncatus*)<br>(Benke, 1993) | Spectacled porpoise (*Phocoena dioptrica*) |  |
| --- | --- | --- | --- | --- | --- |
|  | KS20-20Lc |  |  | KS20-07Pd | KS15-29Pd<br>/VT3347 |
| Side (Right and Left) | R | L | R | R | R |
| FLB (flipper width) | 111 | 111 | 77 | 88.56 | 70.6 |
| FLL (flipper length) | 342 | 335 | n/a | 221.5 | 165 |
| ML (manus length) | 233 | 229 | 70 | 115.4 | 84.6 |
| RL (radius length) | 73.1 | 70.8 | 75 | 62.31 | 48.4 |
| RUD (radius and ulna distal width) | 80.3 | 77.6 | 70 | 49.89 | 42.8 |
| RUP (radius and ulna proximal width) | 63.4 | 64.2 | 64 | 36.4 | 38.6 |
| UL (ulna length) | 65.7 | 64.8 | 66 | 49.77 | 40.2 |

#### Tympano-periotic complex

In hourglass dolphin KS20-20Lc, the right and left tympano-periotic complex weighed 18 and 17 g, respectively (Figure 8). For a full reporting of measurements, refer to Table 6.

**Figure 8:**
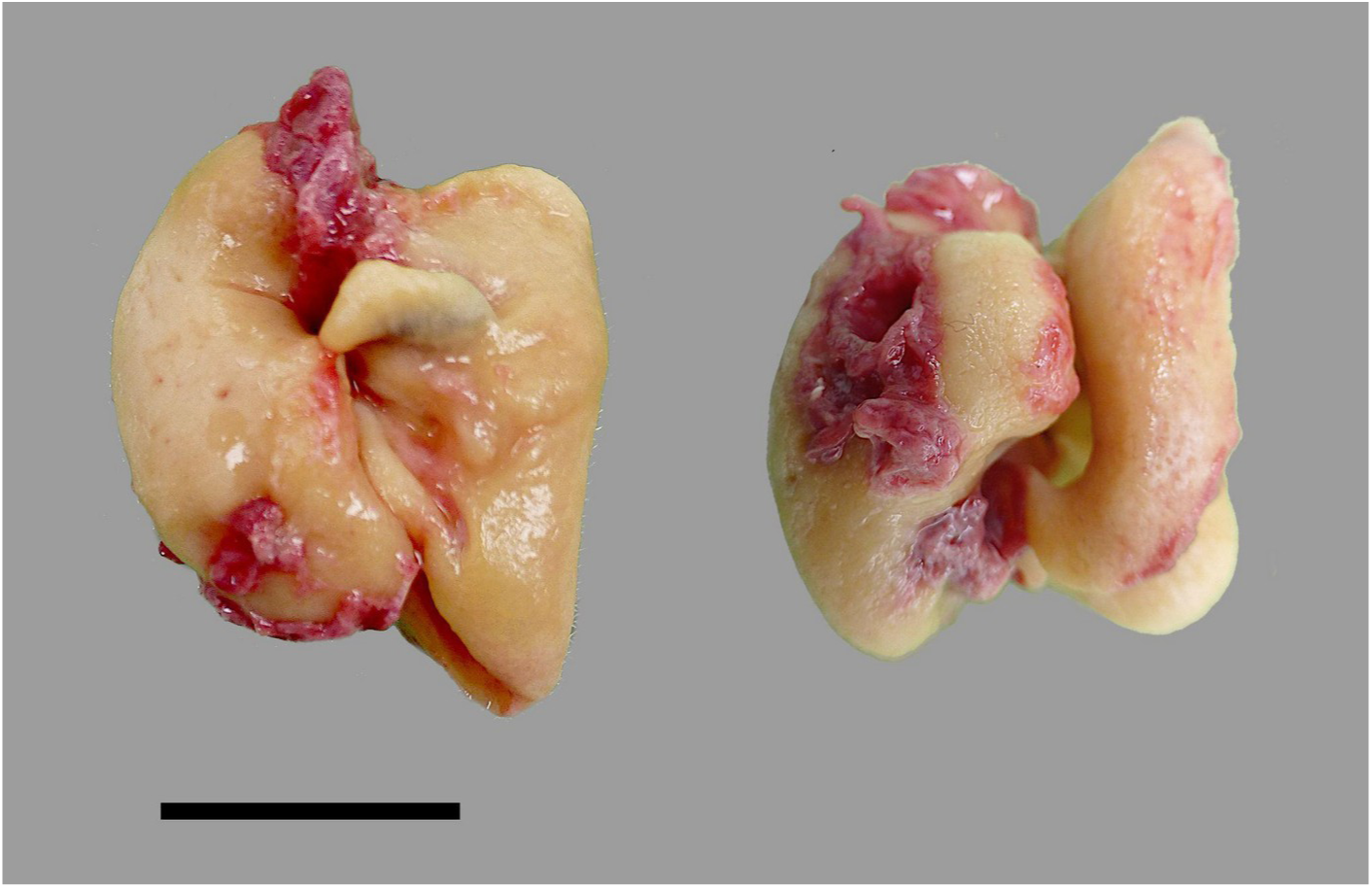
Tympano-periotic complex of hourglass dolphin KS20-20Lc. Scale bar = 1 cm.

**Table 6.** Morphometry of the tympanic-periotic complex. TPC, tympanic-periotic complex; T, tympanic bone; P, periotic bone; L left; R, right. Measurements are shown as length x width x depth.

| Parameter | Hourglass dolphin<br>(*Lagenorhynchus cruciger*) |  | Spectacled porpoise<br>(*Phocoena dioptrica*) |  |  |
| --- | --- | --- | --- | --- | --- |
|  | KS10-28Lc | KS20-20Lc | KS14-45Pd/<br>X2020.76 | KS14-37Pd/<br>X2020.77 | KS20-07Pd |
| T R measurements (mm) | 35 x 21 | 35 x 19 | 32 x 15 | 32 x 18 | 30 x 18 |
| T L measurements (mm) | 34 x 18 | 34 x 19 | 31 x 12 | 31 x 15 | 31 x 19 |
| P R measurements (mm) | 30 x 22 | 29 x 15 x 8 | 36 x 19 | 31 x 15 | 32 x 18 |
| P L measurements (mm) | 31 x 20 | 29 x 18 x 12 | 36 x 20 | 32 x 20 | 31 x 20 |

### Visceral anatomy

#### Respiratory system

In both species we found robust tracheal cartilaginous rings, visible bronchial cartilages down to the smallest distinguishable airway, a right tracheal bronchus slightly cranial to the primary bronchial division and no apparent lobation of either the left or right lung. Some parameters can be found in Table 7. In hourglass dolphins, both the lungs terminated in alignment with rib 12. In spectacled porpoises, the left lung terminated approximately at rib 12, with the right lung aligned with rib 10 (Figures 9 and 10). Histologically, in both species it was possible to observe presumed myoelastic sphincters surrounded by cartilage in the smaller bronchioles (Figure 11).

**Figure 9:**
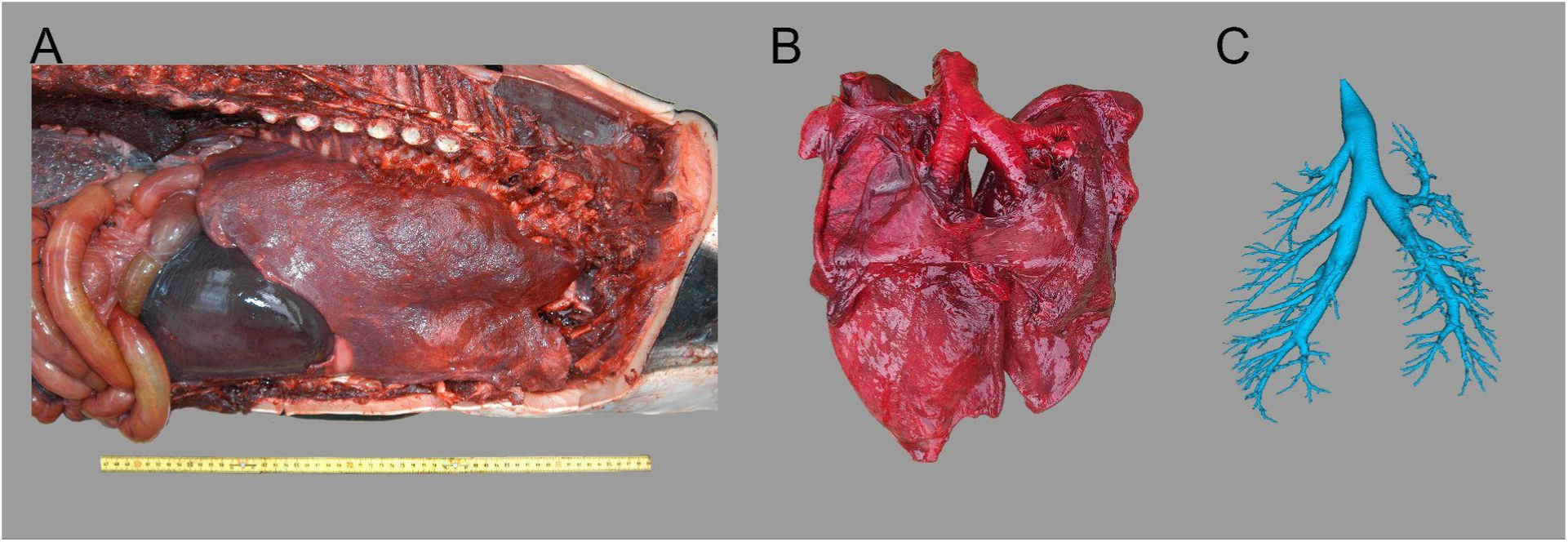
Thorax with lungs in their topographical location without ribs (A), isolated dorsal and ventral aspects (B), and as a 3D reconstruction, showing the trachea and bronchial tree of hourglass dolphin KS10-28Lc. Scale on the left is common to all images.

**Figure 10.**
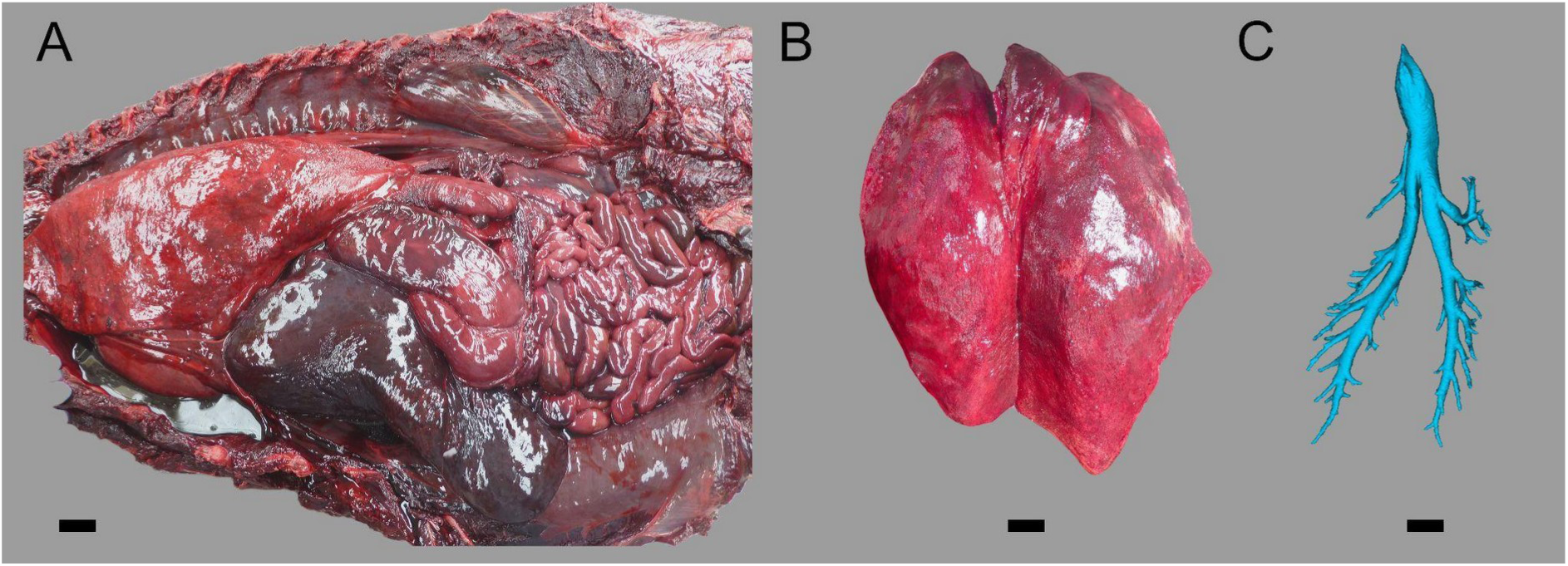
Representation of spectacled porpoise KS20-07Pd lungs (A) in their topographical location and (B) after removal. (C) 3D reconstruction of the trachea and bronchial tree of spectacled porpoise KS20-07Pd. Scale bar = 2 cm.

**Figure 11.**
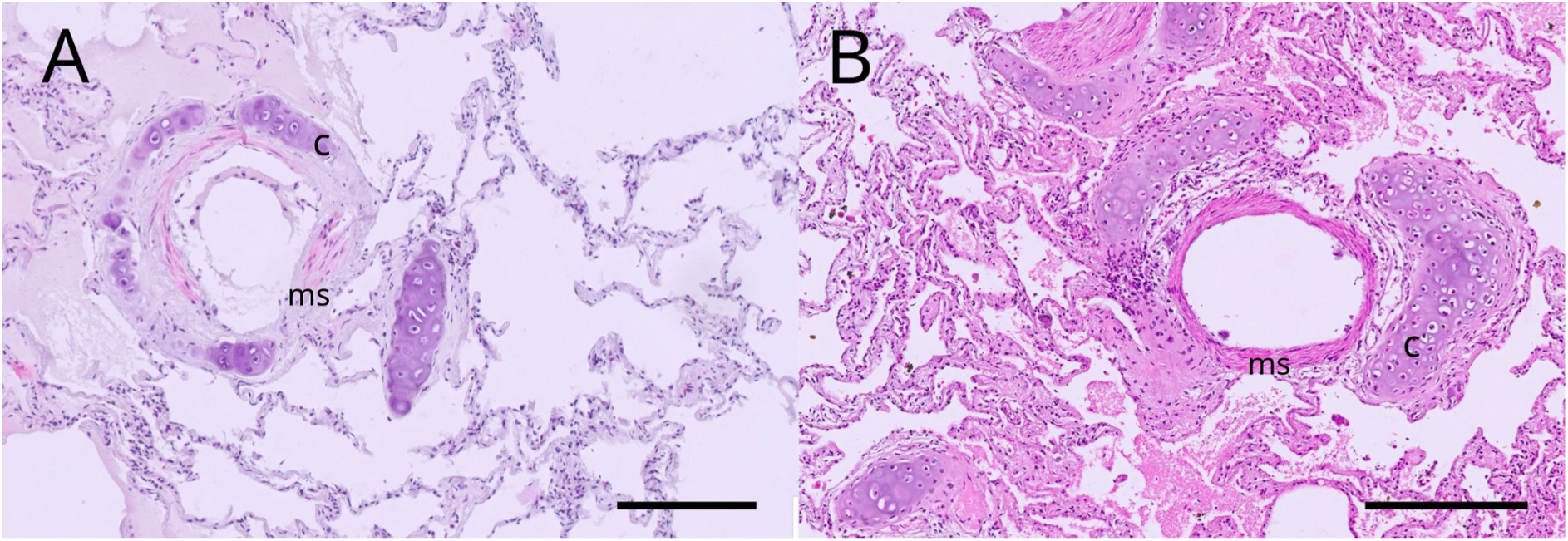
Photomicrograph of the (A) hourglass dolphin KS20-20Lc and (B) spectacled porpoise KS20-07Pd small bronchioles showing the presumed myoelastic sphincters (ms) surrounded by cartilage (c). Scale bar = 100 μm. Hematoxylin-eosin stain.

**Table 7:** Lung parameters. Measurements are shown as (craniocaudal) length x (dorsoventral) height x (lateromedial) width.

| Parameter | Hourglass dolphin<br>(*Lagenorhynchus cruciger*) | Spectacled porpoise (*Phocoena dioptrica*) |  |  |
| --- | --- | --- | --- | --- |
|  | KS20-20Lc | KS14-37Pd/<br>X2020.77 | KS15-29Pd/<br>VT3347 | KS14-45Pd/<br>X2020.76 |
| Total weight (g) | 3100 | 600,3 (L); 673 (R) | 328,2 (L); 365.35 (R) | 1456 (L);<br>1078 (R) |
| Total volume (L) | 3.4 | n/a | n/a | n/a |
| Right lung<br>measurements (mm) | 440x190x98 | 400x120 | n/a | n/a |
| Left lung<br>measurements (mm) | 420x170x75 | 400x130 | n/a | n/a |

#### Circulatory and lymphatic systems

##### Heart and vessels

In both species, the heart was relatively flat dorsoventrally and triangular in shape, with wide and flat auricles. The left ventricle was larger, with a thicker wall compared to the right. Table 8 summarises the most important measurements in some specimens.

**Table 8:** Heart parameters. Measurements shown as width x length x diameter.

| Parameter | Hourglass dolphin<br>(*Lagenorhynchus cruciger*) | Spectacled porpoise (*Phocoena dioptrica*) |  |  |
| --- | --- | --- | --- | --- |
|  | KS20-20Lc | KS14-37Pd/<br>X2020.77 | KS15-29Pd/<br>VT3347 | KS14-45Pd/<br>X2020.76 |
| Weight (g) | 296.6 | 562.9 | 269.6 | 1037 (approx.<br>vol. of 1857 cm<sup>3</sup>) |
| Measurements (mm) | n/a | 180x220 | n/a | 192x210x45.6 |

The heart of both hourglass dolphins was located between the intercostal spaces 1 to 5, lying on the sternum with its major axis oriented laterally, so that the right and left ventricles aligned to the right sides, respectively (Figure 12A, B). The paraconal groove was relatively deep with large arteries covered in fat. The diameter of the aorta was approximately 3 cm.

**Figure 12.**
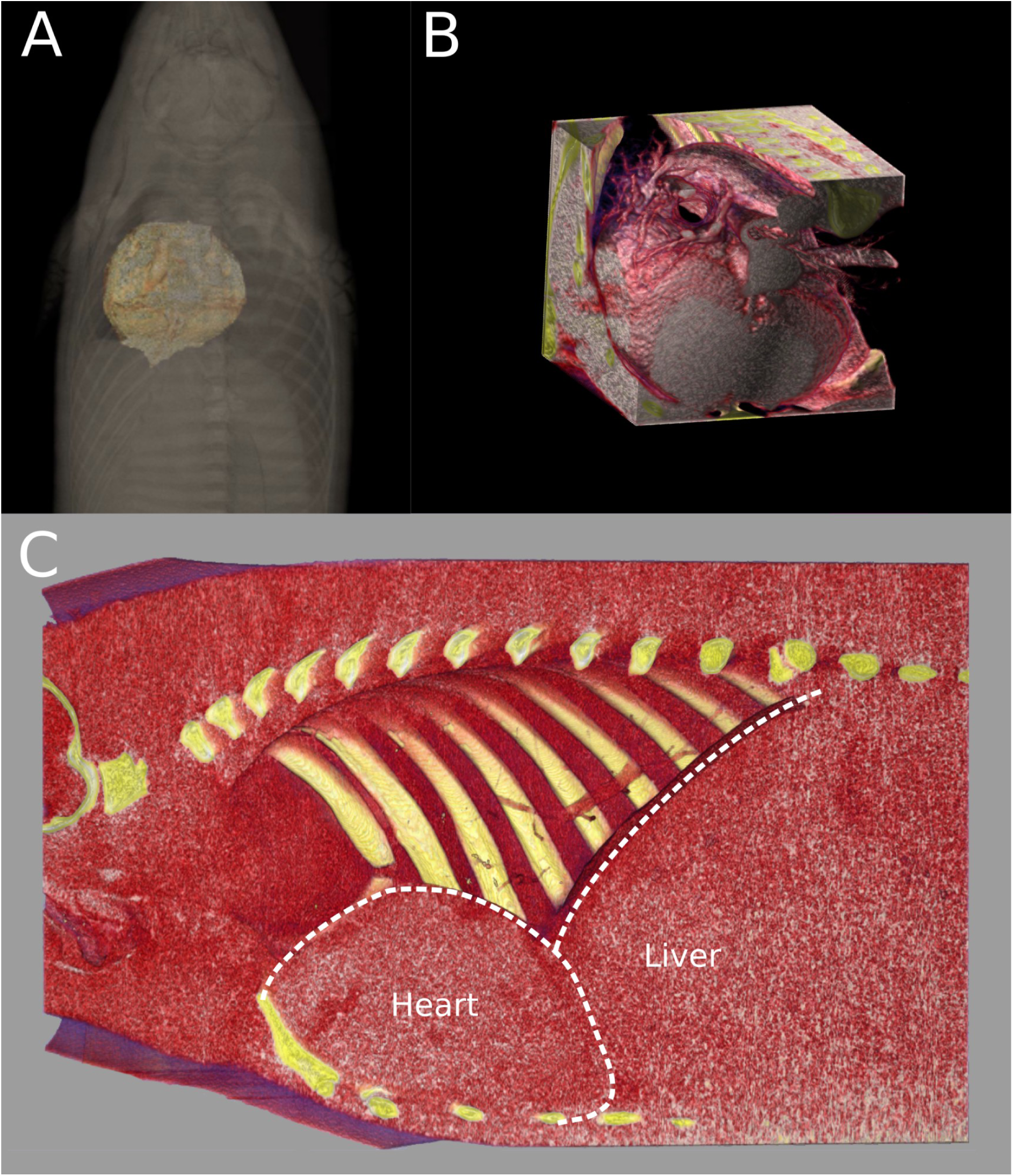
3D reconstruction of the heart in (A, B) hourglass dolphin KS20-20Lc and (C) spectacled porpoise KS20-07Pd. (A) views the heart dorsally, showing its topography in the thoracic cavity, (B) illustrates the heart in isolation, with the aortic arch and onset of aorta shown and (C) presents a 3D model of the heart, sectioned along the sagittal plane, revealing detailed views of the right side of the heart and the liver.

The hearts of spectacled porpoises KS20-07Pd and KS14-45Pd/X2020.76 were located on the sternum between intercostal space 1 and 4 (Figure 12C), and had an approximate volume of 1320 cm^3^. The heart of KS20-07Pd weighed 813 g, with an aorta diameter of 3.9 cm recorded. Several vertebral arteries extended from the heart to the thoracic rete mirabile along its course in the thorax.

In spectacled porpoise KS14-45Pd/X2020.76, it was also possible to remove the dorsal fin and scan it independently (Figure 13). The images displayed the characteristic pattern of the countercurrent exchange system in cetacean appendages, featuring central arteries located along the midline of the dorsal fin and branching dorsally, encircled by circumferential veins (Figure 13). We hypothesised that the central arteries originating from the body were represented by the hypodense areas in the CT scan, as these structures possessed rigid walls and may have lost their blood content, resulting in air-filled cavities while maintaining their structural integrity. However, instead of observing a singular line of arteries, we identified multiple lines, exhibiting an inhomogeneous distribution (Figure 13C). The 3D rendering revealed large vessels only in the central section of the dorsal fin, with scarcity in the rostral and caudal areas. Bifurcation varied, with some vessels bifurcating at the base while other vessels bifurcated more toward the tip, and others at mid-height (Figure 13D). Cranial and caudal arteries tended to curve cranially and caudally at the tip of the fin, respectively, while central arteries ran perpendicular to the fin.

**Fig 13.**
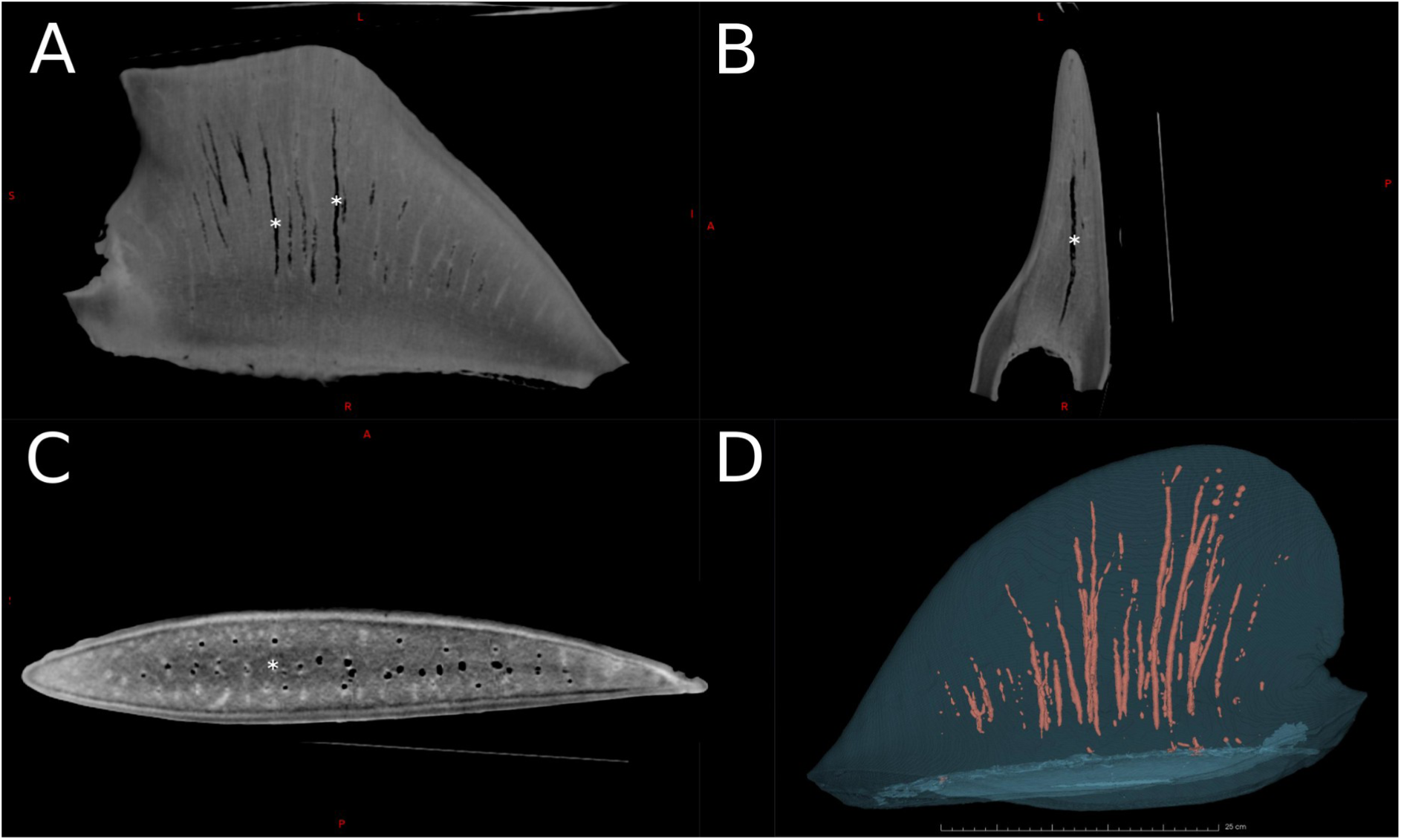
CT scan of the dorsal fin of spectacled porpoise KS14-45Pd/X2020.76. (A), (B) and (C) show the sagittal, transversal and axial planes at its base, respectively. (D) shows the 3D reconstruction with the render of the arteries. In (A), (B) and (C), the asterisks highlight the central arteries. These are shown as hypodense (dark) regions. The circumferential veins surrounding the arteries, due to their weaker wall and size, may have collapsed, thus appearing as small hyperdense areas. Conversely, the peripheral veins were visible as hyperdense areas.

#### Spleen

In both specimens the spleen was found, with main parameters provided in Table 9. In spectacled porpoise KS20-07Pd, an accessory spleen was further recorded (Figure 14). In hourglass dolphin KS20-20Lc, prescapular lymph node weights and measurements were 15 g (L) and 20 g (R) in and 51 x 40 x 11 mm (L) and 72 x 39 x 15 (R), respectively. Histologically, in both species the spleen was composed of an external capsule which also sent trabecule into the underlying parenchyma, a white pulp composed of immune cells distributed in the red pulp (Figure 15). Mesenteric lymph nodes were present at the root of the mesentery.

**Figure 14.**
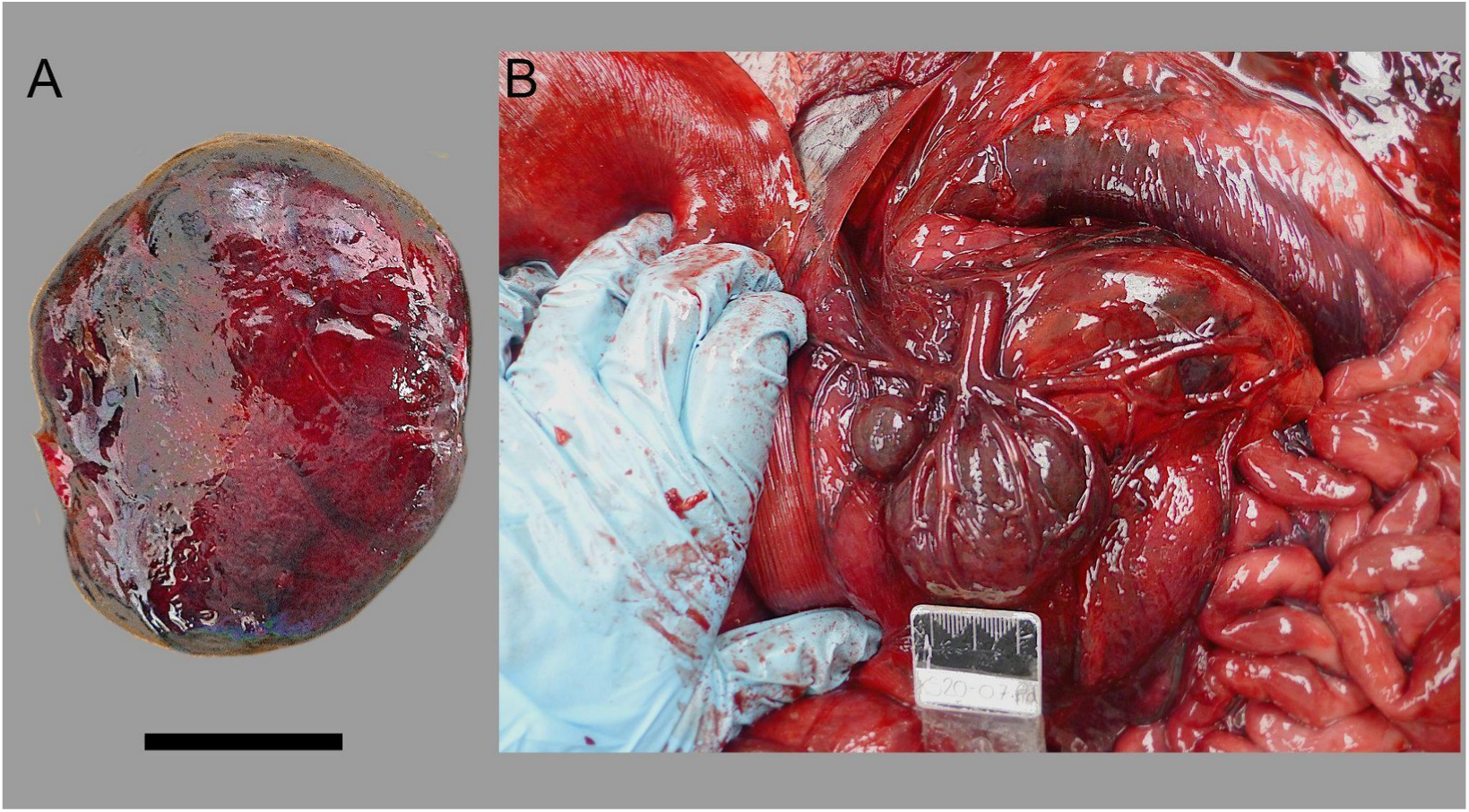
Spleen of porpoise KS20-07Pd (A), with presence of accessory spleen (B) shown to the left of the primary spleen. Scale bar = 2 cm.

**Figure 15.**
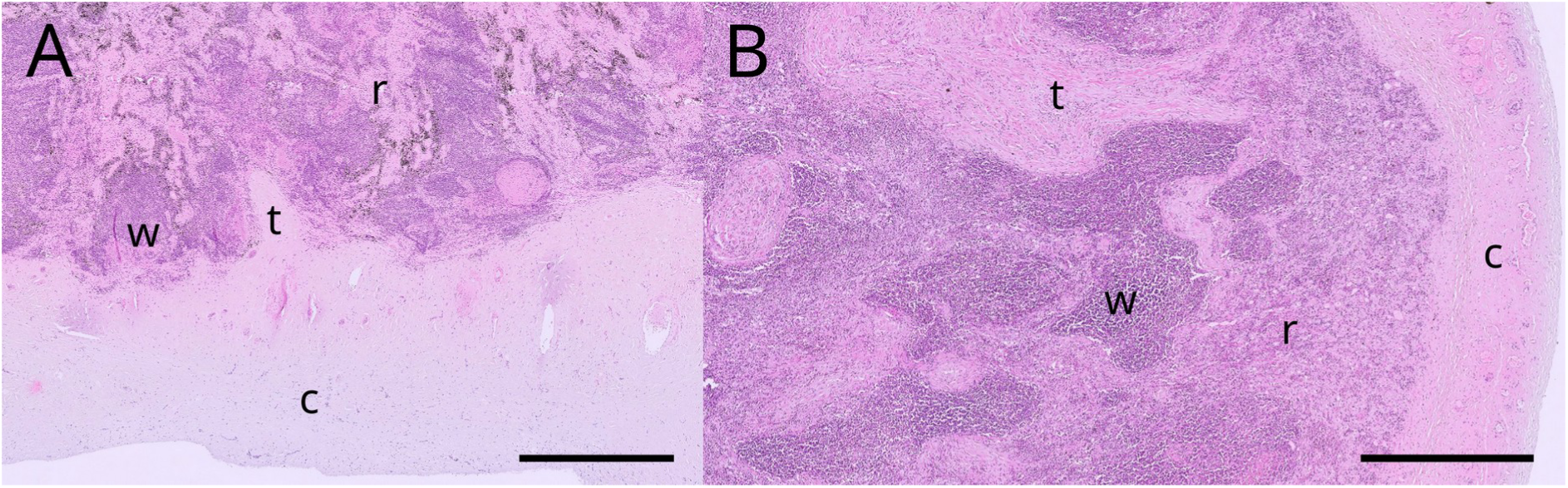
Photomicrograph of the (A) hourglass dolphin KS20-20Lc and (B) spectacled porpoise KS20-07Pd spleen. Note the capsule (c) sending trabecule (t) into the parenchyma, in turn divided into white pulp (w) and red pulp (r). Scale bar = 1000 μm. Hematoxylin-eosin stain.

**Table 9.** Spleen parameters. Measurements shown as length x width x diameter (mm).

|  | Hourglass dolphin<br>(*Lagenorhynchus cruciger*) | Spectacled porpoise (*Phocoena dioptrica*) |  |  |
| --- | --- | --- | --- | --- |
|  | KS20-20Lc | KS15-29Pd/<br>VT3347 | KS14-45Pd/<br>X2020.76 | KS20-07Pd |
| Weight (g) | 20 | 19.93 | 16 | 27 |
| Measurements (mm) | 63x40x15 | 40x35 | n/a | 59x48x17 |

#### Digestive system

##### Mouth and upper digestive tract

While most teeth were intact in the hourglass dolphin KS10-28Lc, most teeth in hourglass dolphin KS20-20Lc were worn to the gum (LR: 1-11; LL: 1-13; UR: 1; UL: 1-5 and 9-24). This indicates an older specimen, which was further supported in the mineralization of the pectoral limb (Figure 7) in KS20-20Lc. Teeth on KS15-29Pd were only partially erupted and teeth of KS14-37Pd/X2020.77 were not examined. The dental formula of examined specimens can be viewed in Table 10.

**Table 10.**
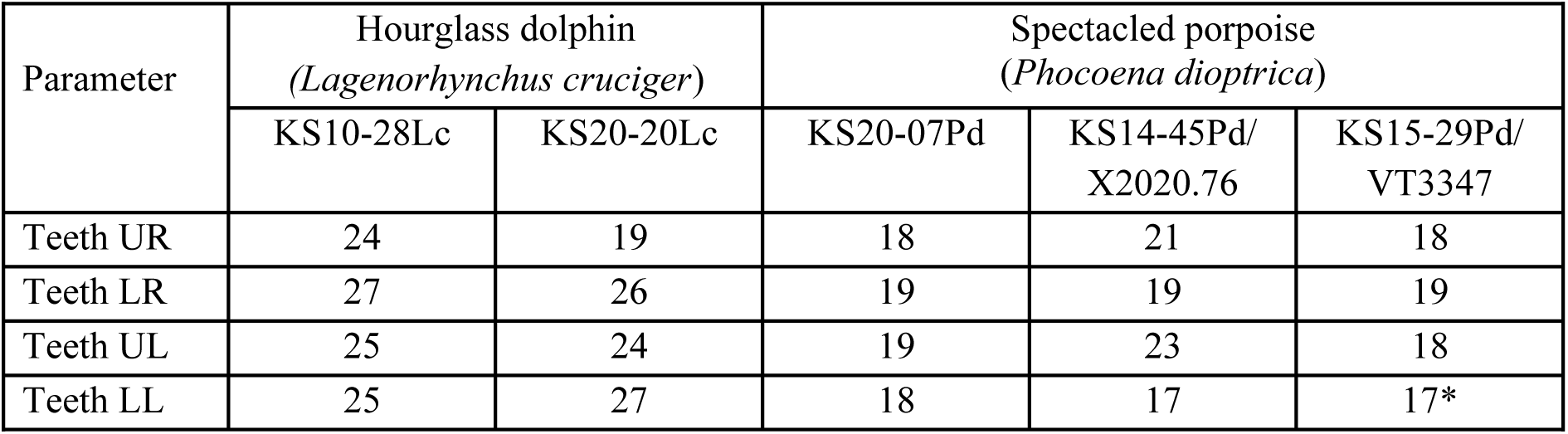
Dental formula of selected specimens examined. Note: UL, upper left; UR, upper right; LL, lower left; LR, lower right.

| Parameter | Hourglass dolphin<br>(*Lagenorhynchus cruciger*) |  | Spectacled porpoise<br>(*Phocoena dioptrica*) |  |  |
| --- | --- | --- | --- | --- | --- |
|  | KS10-28Lc | KS20-20Lc | KS20-07Pd | KS14-45Pd/<br>X2020.76 | KS15-29Pd/<br>VT3347 |
| Teeth UR | 24 | 19 | 18 | 21 | 18 |
| Teeth LR | 27 | 26 | 19 | 19 | 19 |
| Teeth UL | 25 | 24 | 19 | 23 | 18 |
| Teeth LL | 25 | 27 | 18 | 17 | 17* |

The pointed tongue in KS20-20Lc measured 13.3 (L) x 6.3 cm (W), with no anterolateral papillae. Six vallate papillae were present at the root of the tongue, arranged in a V orientated towards the pharynx (arrowhead in Figure 16).

**Figure 16.**
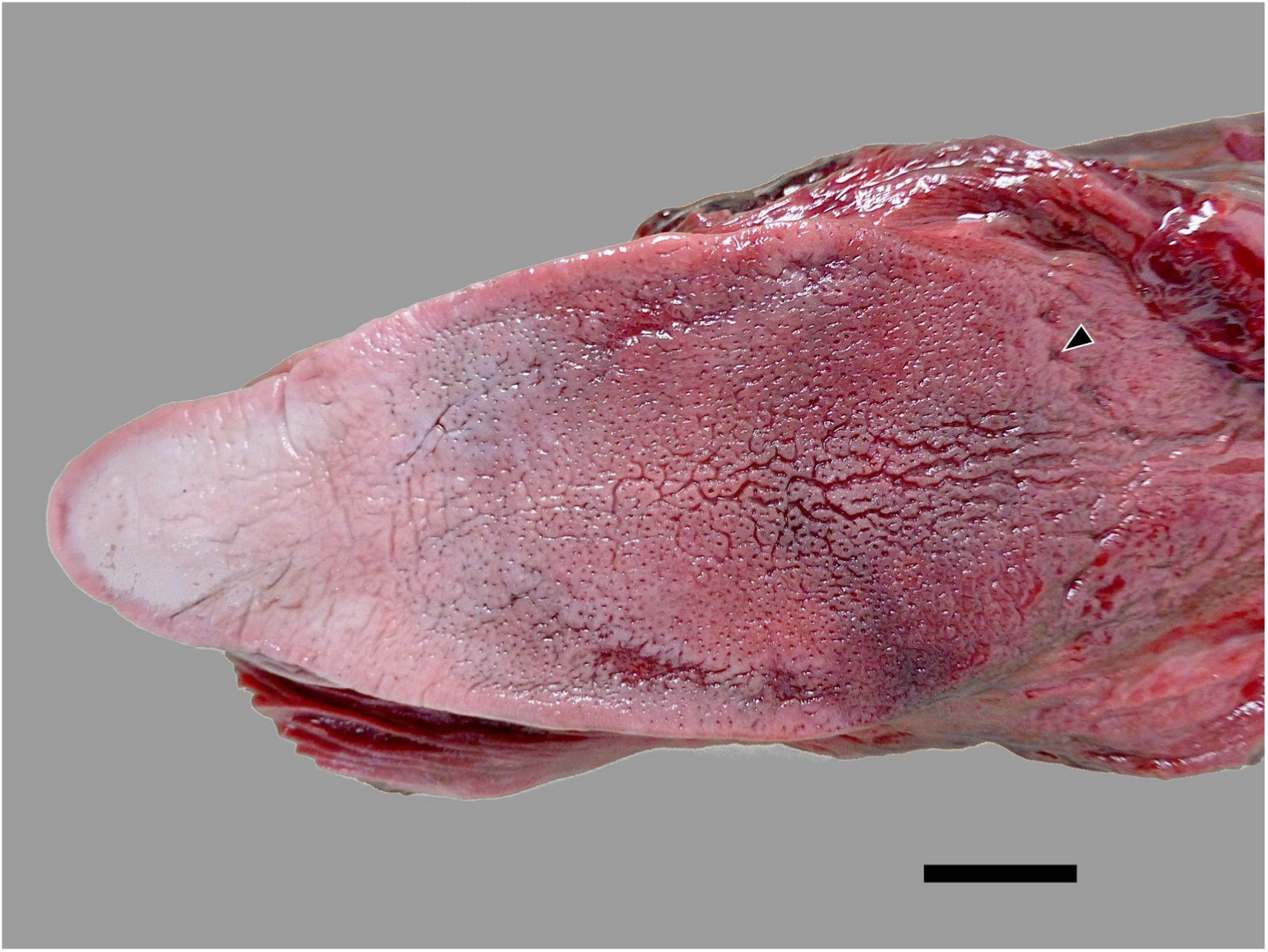
Tongue of the hourglass dolphin KS20-20Lc. Arrowhead denotes vallate papilla in a V orientated towards the pharynx. Scale bar = 2 cm.

#### Stomach complex and intestine

The stomach chambers of all specimens resembled the typical delphinid pattern: one forestomach, one main stomach and one pyloric stomach. Similarly, in both species the intestine was a unique “tube” without macroscopical distinction between small and large parts, and lacking a caecum. The mesenteric lymph nodes were clearly identified (Figure 17). Microscopically, the main stomach (second chamber) mucosa was thick with the typical pattern of other mammals such as the presence of glandular cells secreting mucus and HCl (Figure 18A); the pyloric stomach (third chamber) mucosa was thinner with columnar epithelium and tubular glands (Figure 18B). Finally, the mucosa of the jejunum was characterized by the typical presence of villi (Figure 18C).

**Figure 17.**
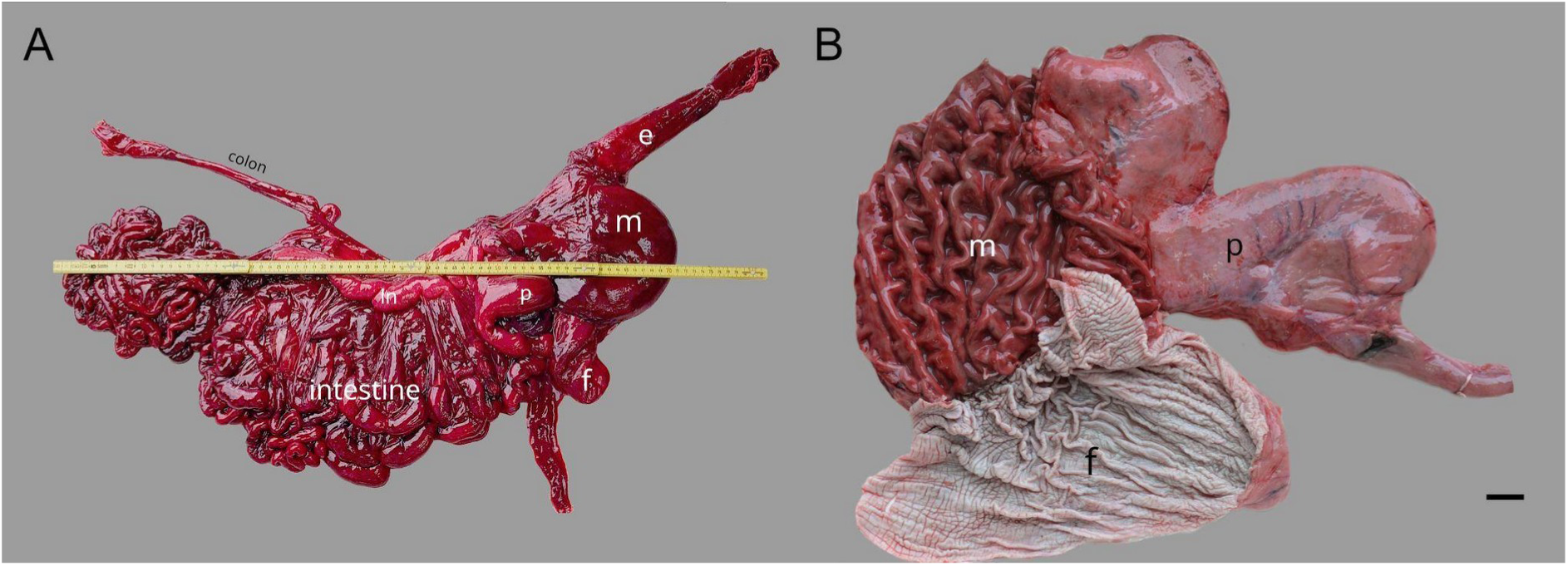
Gastrointestinal components of the digestive system. (A) whole gastrointestinal system of hourglass dolphin KS20-20Lc, starting from the oesophagus and terminating at the rectum. (B) gastric chambers of the spectacled porpoise (KS14-45Pd/X2020.76). e, oesophagus; f, forestomach; i, intestine; ln, mesenteric lymph nodes; m, main stomach; p, pyloric stomach. Scale bar = 1 cm.

**Figure 18.**
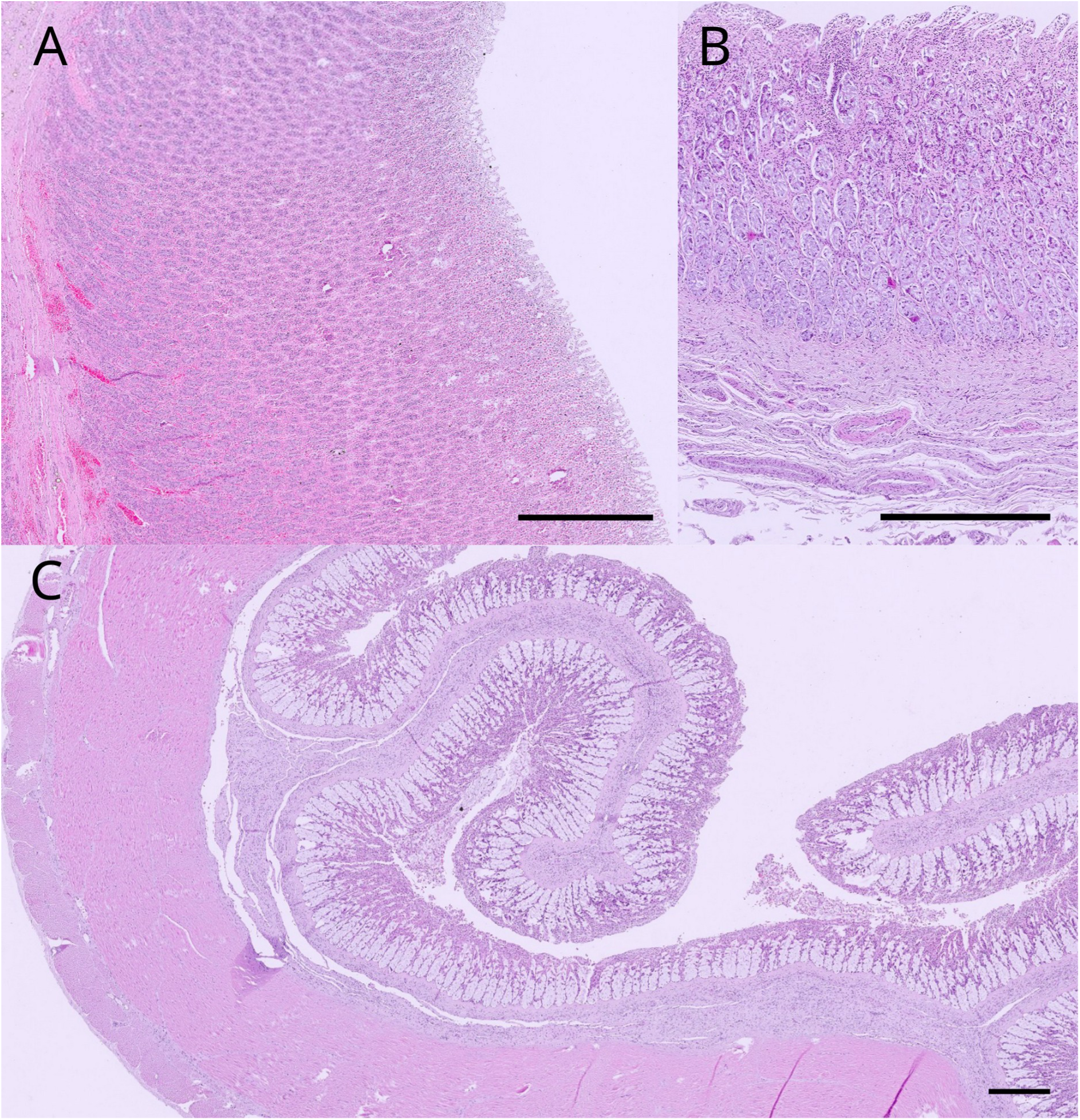
Microphotograph of the different mucosae of (A) porpoise KS20-07Pd main stomach; (B) hourglass dolphin KS20-20Lc pyloric stomach and (C) porpoise KS20-07Pd jejunum. Scale bar = 200 μm. Hematoxylin-eosin stain.

#### Liver

In the two hourglass dolphins, liver position ran from the 10th thoracic to the 2nd lumbar vertebrae, on the ventral half of the abdominal cavity (Figure 19A). In KS10-28Lc, the right lobe extended considerably further than the left, which is consistent with its topography and the position (leftward) of the stomachs. The diaphragmatic surface of the right lobe was expanded cranially and contained most of the mass of the organ. The right lobe was separated from the left by a thin falciform ligament, which terminated in a sheet covering the cranial portion of the stomachs (Figure 19B). This was considerably less evident in the KS20-20Lc, where the liver appeared almost divided in even halves, although the right half was thicker than its left counterpart.

**Figure 19.**
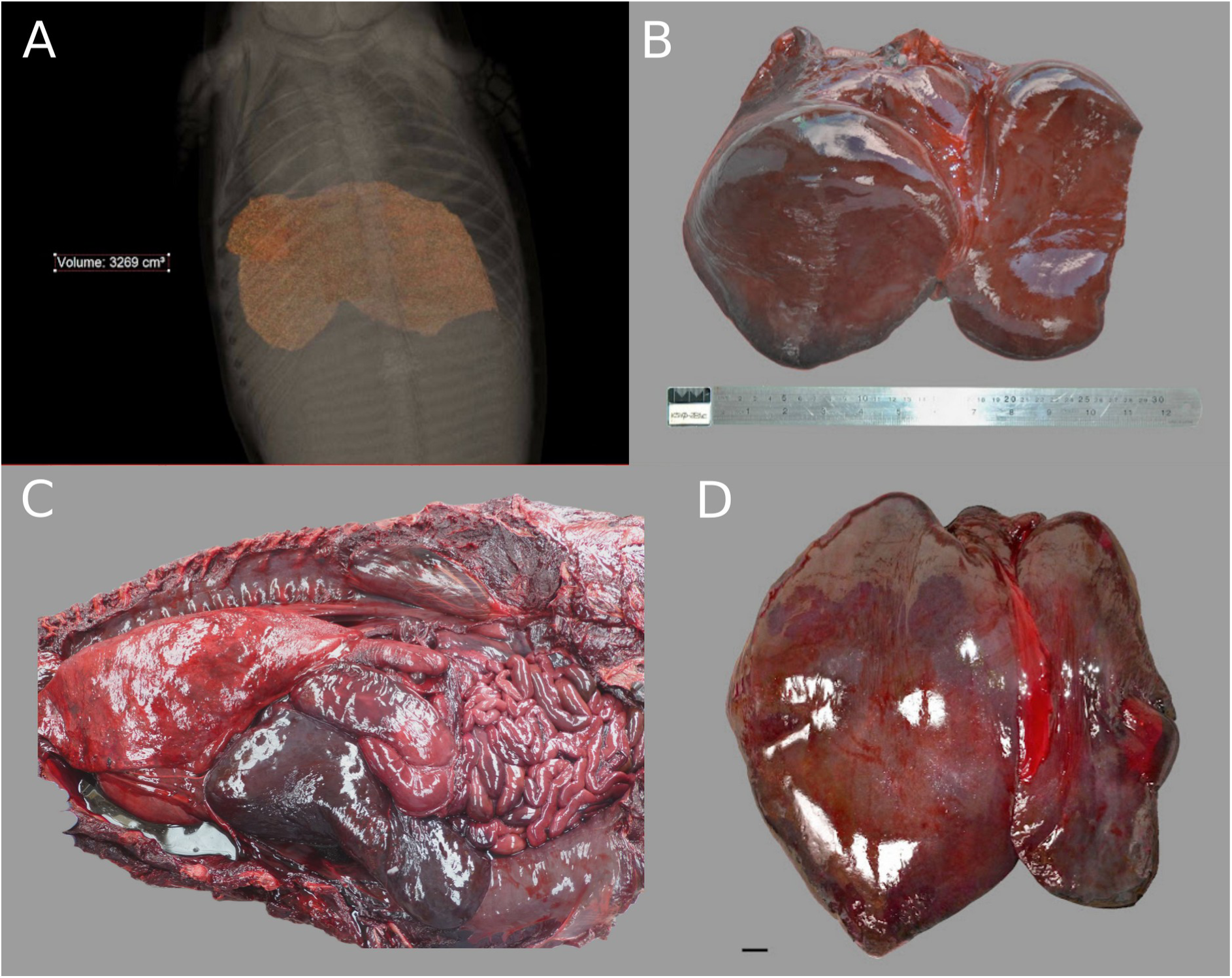
Representation of the liver in the (A, B) hourglass dolphin KS20-20Lc and (C, D) spectacled porpoise KS20-07Pd. (A) and (C) showed the liver in its topographical location within the abdominal cavity while (B) and (D) the extracted and isolated organ. Note the difference between the large right compared to smaller left lobe. Scale bar of (D) = 1 cm.

In the spectacled porpoises, the liver was positioned approximately between thoracic rib 8 to lumbar vertebra 4. A highly developed right compared to more inferior left lobe was noted, without the aforementioned diaphragmatic expansion for the two hourglass dolphin specimens. The falciform ligament still clearly divided the main left and right lobes of the liver (Figure 19C, D). In both species we also could not find any venous sinus, apparent lobulation nor gallbladder.

Histologically, in the hourglass dolphin the hepatic parenchyma was formed of lobules, not easily distinguishable due to the absence of connective septa and clear central veins. However, portal triads, in turn composed of a portal vein, hepatic artery and bile duct were identified (Figure 20). No definite muscular wall surrounding the portal vein was found.

**Figure 20.**
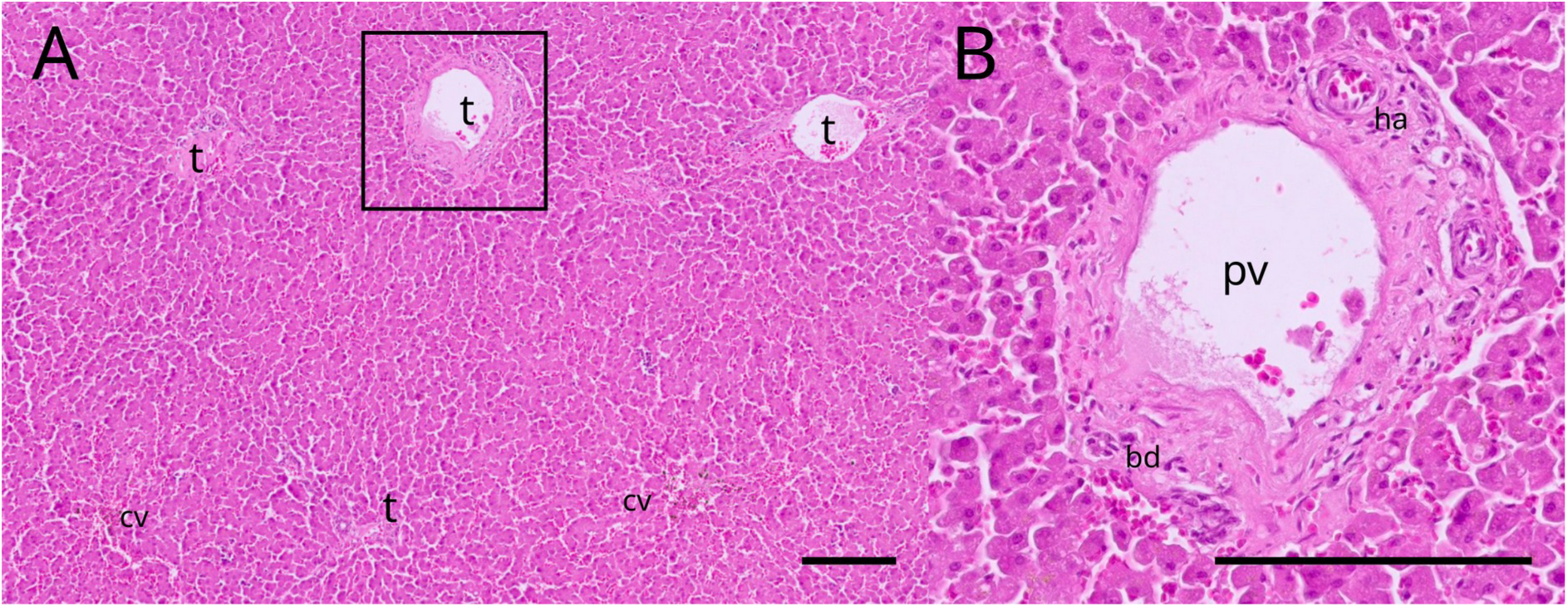
Microphotograph of the KS28-10Lc liver. (A) low-magnification image showing the overall hepatic organization in central veins (cv) surrounded by portal triads (t). (B) high- magnification image of a portal triad of (A) showing the portal vein (pv), hepatic artery (ha) and bile duct (bd). Scale bar = 200 μm. Hematoxylin-eosin stain.

Details on the dimensions and weight of the liver for three specimens (KS10-28Lc, KS20-20Lc and KS20-07Pd) are reported in Table 11.

**Table 11.** Liver morphometry in hourglass dolphin and spectacled porpoise. Measurements shown as width x length x height.

| Parameter | Hourglass dolphin<br>(*Lagenorhynchus cruciger*) |  | Spectacled porpoise<br>(*Phocoena dioptrica*) |  |  |  |
| --- | --- | --- | --- | --- | --- | --- |
|  | KS10-28Lc | KS20-20Lc | KS20-07Pd | KS14-45Pd/<br>X2020.76 | KS14-37Pd/<br>X2020.77 | KS15-29Pd/<br>VT3347 |
| Weight (g) | 1865 | 1964 | 3855 | 4143 | 2368 | 861 |
| Measurements (mm) | n/a | 272 x 309 x 42 | n/a | n/a | n/a | n/a |

#### Endocrine system

##### Adrenal glands

In both species, the adrenal glands were composed of a thick cortex (divided into *zona glomerulosa*, *fasciculata and reticularis*), and a thin medulla. The adrenal glands of hourglass dolphins demonstrated a more ovoidal shape compared to the spectacled porpoises, which were more pyramidal in shape. Microscopically, there were more septa in the spectacled porpoise compared to the hourglass dolphin, with the zona *fasciculata* comprising the thickest component (Figure 21). Details on size and weight of the adrenal glands are reported in Table 12.

**Figure 21.**
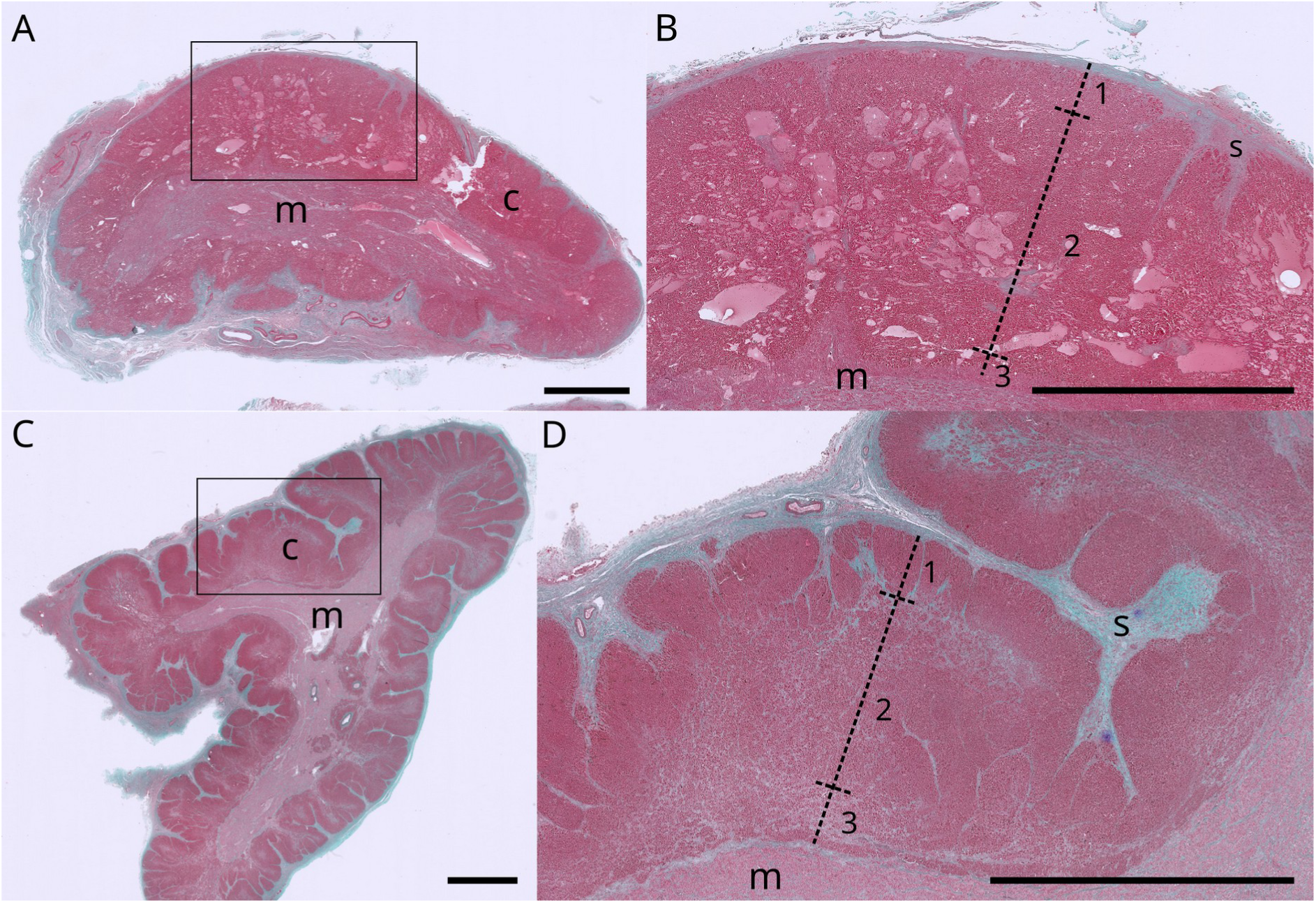
Microphotograph of the adrenal glands of (A, B) hourglass dolphin KS20-20Lc and (C, D) spectacled porpoise KS20-07Pd. (A) and (C) display the whole gland while (B) and (D) offer a subdivision of the cortex into zona *glomerulosa* (1), *fasciculata* (2) and *reticularis* (3). c, cortex; m, medulla; s, septa. Scale bar = 500 μm. Masson’s trichrome stain.

**Table 12.** Adrenal gland morphometry. Measurements shown as length x width x diameter. Note: L = left; R = right.

| Parameter | Hourglass dolphin<br>(*Lagenorhynchus cruciger*) |  | Spectacled porpoise (*Phocoena dioptrica*) |  |  |
| --- | --- | --- | --- | --- | --- |
|  | KS10-28Lc | KS20-20Lc | KS20-07Pd | KS14-37Pd<br>/X2020.77 | KS14-45Pd/<br>X2020.76 |
| Adrenal R weight (g) | n/a | 8 | 7 | 19.2 | 15 |
| Adrenal R measurements (mm) | n/a | 78 x 28 x 9 | 55 x 30 x 17 | 67 x 35 x<br>n/a | n/a |
| Adrenal L weight (g) | n/a | 7 | 8 | 18.4 | 11 |
| Adrenal L measurements (mm) | n/a | 52 x 23 x 7 | 51 x 34 x 10 | 55 x 26 x<br>n/a | n/a |

#### Urogenital system

##### Kidneys

Each kidney in the hourglass dolphins had ca. 300 reniculi, similar to that of bottlenose dolphin *Tursiops truncatus* (Cozzi et al., 2017). Renal structure was typical of cetaceans, with the muscular basked (*sporta perimedullaris*) and renicular arterioles dividing the cortex and medulla (Figure 22). Details on the morphometry of the kidneys are further summarised in Table 13.

**Figure 22.**
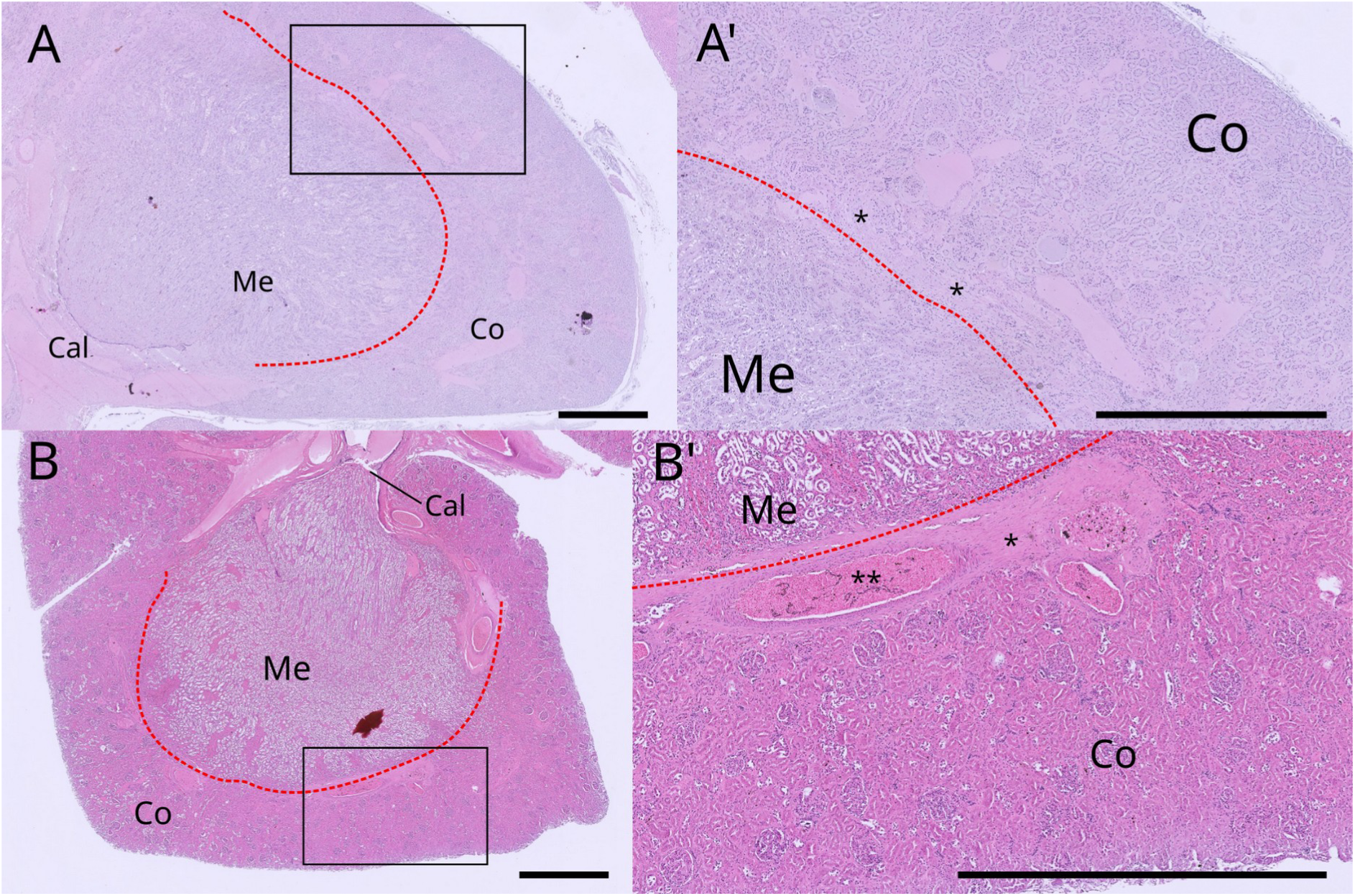
Microphotograph of reniculus of (A, A’) hourglass dolphin KS20-20Lc and (B, B’) spectacled porpoise KS20-07Pd. (A) and (B) show the whole structure with the red dotted line dividing the cortex (Co) from the medulla (Me), which in turn collects the urine in the calyx (cal). (A’) and (B’) show the muscular basket (*) and renicular arterioles (**). Scale bar = 1 cm. Hematoxylin-eosin stain.

**Table 13.**
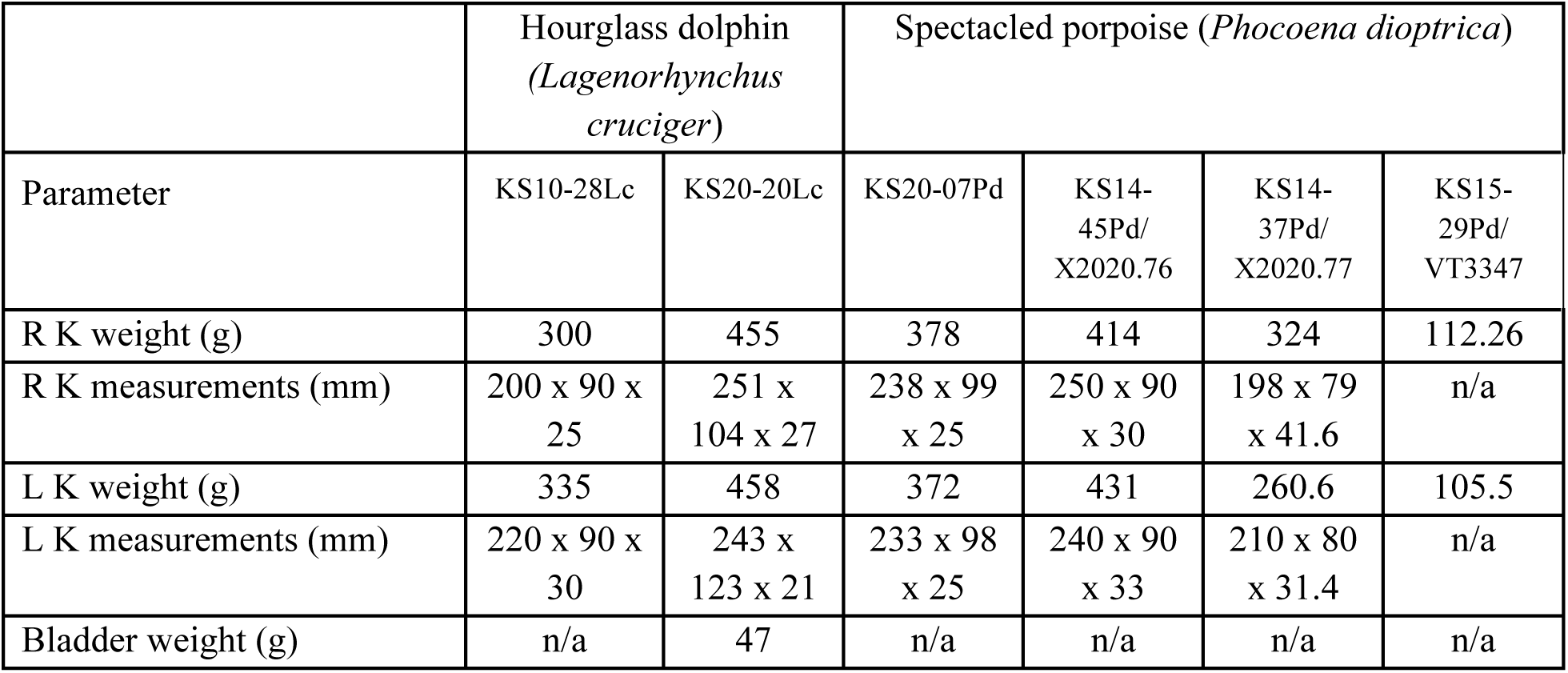
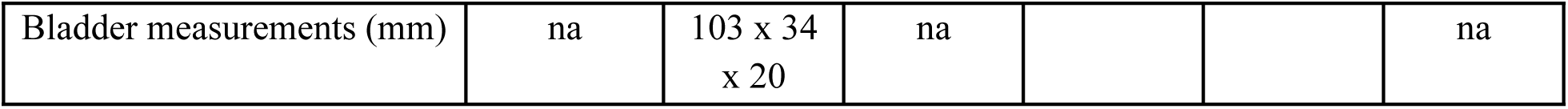
Morphometry of the kidneys. K, kidney; L, left; R, right. Measurements: length x width x diameter.

#### Gonads

In KS15-29Pd/VT3347, the left ovary weighed 1.9 g, was 15 mm in length, 10 mm in width and 5 mm in diameter. The uterus was 30 mm in length and 15 mm in width, belonging to an immature female.

In males of both species, the penis was S-shaped and displayed the retractor muscles. The apex was thin and the prostate gland clear (Figure 23).

**Figure 23.**
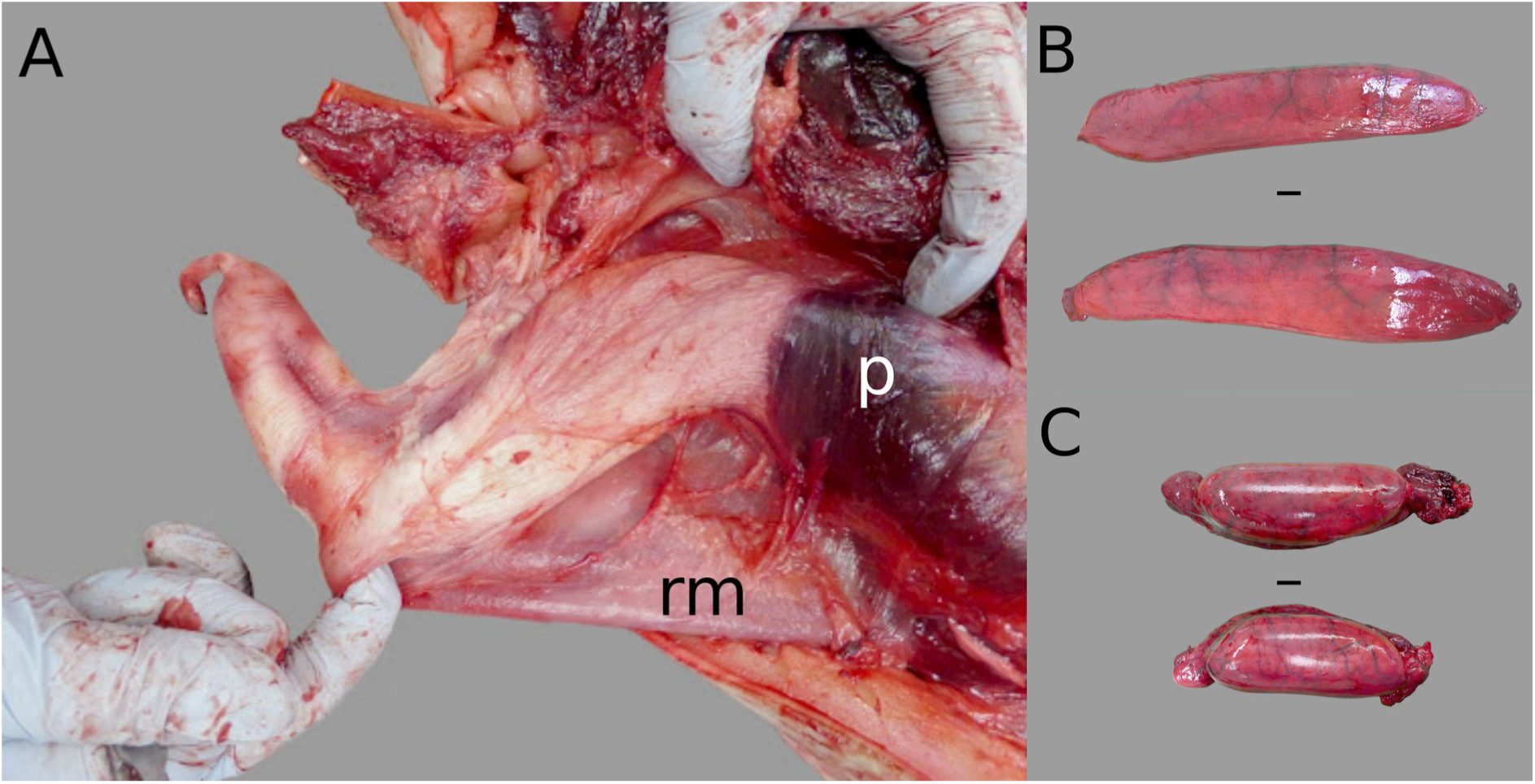
Male reproductive system: (A) Penis of hourglass dolphin KS20-20Lc with retractor muscle (rm) and prostate (p) covered by the muscles ischiocavernosus and bulbo-urethralis; (B) testes of hourglass dolphin KS20-20Lc; (C) testes of spectacled porpoise KS20-07Pd. Scale bar = 1 cm.

In the five males, testes were large, as usual in adult cetaceans, with a white-pearl colour, and an epididymis running on the dorsal border of the testicle for most of its length. However, the testes were more elongated in the hourglass dolphins compared to the spectacled porpoise. The size of the testes in both species indicated that all males were sexually mature. In the case of KS20-20Lc, sexual maturity was confirmed by the presence of spermatozoa on histological examination of testis.

Details on the morphometry of the genital apparatus are reported in Table 14.

**Table 14.** Morphometry of the male genital apparatus. Note: (T) testis, (E) epididymis; L, left; R, right. Note: Measurements are reported as length x width x diameter (mm).

| Parameter | Hourglass dolphin<br>(*Lagenorhynchus cruciger*) |  | Spectacled porpoise<br>(*Phocoena dioptrica*) |  |  |
| --- | --- | --- | --- | --- | --- |
|  | KS10-28Lc | KS20-20Lc | KS20-07Pd | KS14-45Pd<br>/X2020.76 | KS14-37Pd<br>/X2020.77 |
| R T + E weight (g) | 315 | 434 | 323 | n/a | 119.9 |
| R T - E weight (g) | 267 | 369 | 276 | n/a | n/a |
| R T measurements -<br>length x width x depth<br>(mm) | 305 x 59 | 350 x 72 x<br>21 | 161 x 68 x<br>39 | 131 x 51 x<br>n/a | 150 x 50 x<br>n/a |
| L T + E weight (g) | 360 | 466 | 285 | n/a | 124.5 |
| L T - E weight (g) | 270 | 383 | 249 | n/a | 82.2 |
| L T measurements -<br>length x width x depth<br>(mm) | 285 x 55 | 375 x 74 x<br>19 | 152 x 61 x<br>39 | 138 x 50 x<br>n/a | 150 x 50 x<br>n/a |
| Postanal hump<br>measurements (cm) | 4.3 x 28 | 6.5 x 5.2 x<br>28 | n/a | n/a | n/a |
| Penis length (mm) | n/a | 300 | n/a | ~190 | n/a |

## Discussion

The anatomical description of little known or endemic species is rarely for the sole purpose of describing a new species, but rather to offer insight on where the species fits taxonomically. Here, we provided an anatomical overview and description of two seldom reported species, the hourglass dolphin and the spectacled porpoise, using not only photography and conventional histology but also computed tomography including three-dimensional reconstructions. This work was only possible thanks to the collaborative efforts of many individuals spanning multiple teams within and beyond Aotearoa, New Zealand.

In terms of size, both species were within similar ranges of total body length, although male hourglass dolphins weighed marginally less than what was reported for adult males (∼90 kg). Skeletal features, including condylo-basal length and dental characteristics, aligned with previously reported ranges for these species (Goodall et al., 1997; Brownell, 1999; Brownell and Donahue, 1999; Gazitúa et al., 1999; Evans et al., 2001; Pinedo, 2002; Fernández et al., 2003). The anatomical features of other organs, such as the spleen, gastrointestinal tract, liver, adrenal glands, kidneys, and male reproductive systems, were similar to those of other odontocetes. This similarity reinforces the idea of a shared evolutionary lineage and common functional adaptations among marine mammals (Cozzi et al., 2017). This included the characteristic ‘keel’ of the hourglass dolphin, which, as in other delphinids, is composed of blubber and dense connective tissue.

The respiratory anatomy revealed a lack of lobation and the presence of a right tracheal bronchus, consistent with findings in other cetaceans (Fanning and Harrison, 1974; Cozzi et al., 2017). Although systematic measurement of lung weight and total lung capacity was challenging, qualitative evaluation indicated that both species possess relatively large lungs for their body size. This observation aligns with the “short dive, big lung” relationship observed by Piscitelli et al. (2010, 2013), suggesting that their lung size is adapted for this diving behaviour. Meanwhile, observation of the cardiocirculatory system revealed that, in terms of topography and anatomy, the heart in both species aligns with that of other cetaceans (Cozzi et al., 2017).

The dorsal fin of the spectacled porpoise presented unique characteristics, including its size and blood supply. The dorsal fin of other cetaceans studied in captivity has been demonstrated to be a heterogeneous thermoregulatory window, together with the fluke and flippers (Meagher et al., 2002; Cozzi et al., 2017; Plön et al., 2018; Favilla et al., 2023), allowing these animals to conserve or dissipate body heat as required. In particular, vascularisation of the dorsal fin is hypothesised to cool down the male gonads (Plön et al., 2018). Our observations revealed large vessels branching extensively throughout the dorsal fin, reaching up to the tip. These findings were consistent with those reported by Plön and colleagues (2018) in their studies of the Indo-Pacific humpback dolphin (*Sousa plumbea*) and the Indo-Pacific bottlenose dolphin (*Tursiops aduncus*). However, we noted an inexplicable paucity of vessels in the cranial and caudal regions of the fin. This inconsistency may be due to technical limitations in detecting these vessels on CT or a true absence, which would require further investigation, including differences between sexes and potentially utilising MRI with contrast agents to enhance the quality of results. Interestingly, it is worth noting that the Dall’s porpoise (*Phocoenoides dalli*), a relative of the spectacled porpoise that also inhabits cold waters, possesses both a small dorsal fin and fluke, in addition to having a relatively thin blubber layer (Jefferson, 2018). This comparison highlights the variability in thermoregulatory strategies among cetaceans, indicating that thermoregulatory mechanisms need further investigation. The large size of the dorsal fin of the spectacled porpoise could therefore, serve multiple functions. A large dorsal fin could minimise lateral torsion during swimming, enhancing propulsive efficiency, while also maximising exposure to sunlight, which can aid in thermoregulation by warming the blood flowing through its extensive vascular network. To a lesser degree, the dorsal fin of the hourglass dolphin was also large with a wide surface area. However, no similar large vessels were observed in the available CT images, unlike the spectacled porpoise. An alternative hypothesis is that the enlarged dorsal fins of males would serve as a marker of sexual maturity. Collectively, these characteristics highlight the roles in which dorsal fins may affect the ecology and survival of the spectacled porpoise.

Regardless of insights provided, our study faced several limitations regarding the opportunistic methodologies employed. For example, not all specimens were dissected in the same manner or imaged uniformly. Variation in CT scan quality arose from access to different machines, and the advance of technology across the ten years of specimen acquisition. More recent scans showed improved resolution compared to the earlier scans. The unattached large dorsal fin of spectacled porpoise KS14-45Pd/X2020.76 hindered complete scans of the entire animal, making it difficult to count the lumbar vertebrae. Additionally, histological analyses were limited to only a selection of specimens, highlighting the need for greater consistency in future examinations of these rare and elusive species. Despite such challenges, the findings presented here support existing literature, while documenting for the first time the vascular pattern of the dorsal fin of the spectacled porpoise.

In conclusion, this anatomical study of the hourglass dolphin and spectacled porpoise underscores their adaptations to marine environments, reflecting evolutionary pressures that shaped their morphology and physiology. The ecology of these species is essentially a blank canvas. Further research is encouraged on their ecological roles and the potential impacts of environmental changes on their populations.

## Acknowledgements

The authors thank all rūnaka including Wairewa, Arowhenua, Onuku, Ōtākou, for allowing access to their taonga and for collaborating on this kaupapa. Special thanks are extended to kaimahi representing the Rūnaka of Ōraka-Aparima, including Riki Dallas and Iain MacCallum and to Te Kauika Tangaroa Charitable Trust for their support of this Kaupapa. We also acknowledge Melanie Young, Joseph Roberts, Robin Smith, Mike Ogle and Jim Fyfe at the Department of Conservation Te Papa Atawhai for facilitating aspects for the cadaver dissection. Finally, we gratefully thank staff and postgraduates of the Cetacean Ecology Research Group (CERG) and School of Veterinary Sciences (SVS) Massey University, the University of Otago, Tūhura Otago Museum, colleagues at the University of Padova, Italy for their logistical support with dissections and histological assessment of tissues, respectively. This paper is dedicated to the late Prof R. Ewan Fordyce FRSNZ (1953- 2023).

## Ethical statement

All sampling was undertaken under the research permits 39239-MAR, RNW/22/2003/182, and RNW/HO/2008/03 (Massey University), 39645-MAR and 48740-MAR (University of Otago) and 39400-MAR (Tūhura Otago Museum) issued by the Department of Conservation Te Papa Atawhai. All research was done with permission from local rūnanga and rūnaka (local indigenous tribal authorities) alongside Department of Conservation Te Papa Atawhai. As no live animals were sampled, we did not require animal ethics approval for this research. The datasets generated and/or analysed during the current study are available from the corresponding author on reasonable request.

## Data availability statement

The data that support the findings of this study are available on request from the corresponding author. The data are not publicly available due to cultural and ethical restrictions.

## Funding resources

KAS was supported by a Rutherford Discovery Fellowship from the Royal Society Te Aparangi

